# Pathlength-selective, interferometric diffuse correlation spectroscopy

**DOI:** 10.1101/2024.06.21.600096

**Authors:** Mitchell B. Robinson, Marco Renna, Nikola Otic, Olivia S. Kierul, Ailis Muldoon, Maria Angela Franceschini, Stefan A. Carp

## Abstract

Diffuse correlation spectroscopy (DCS) is an optical method that offers non-invasive assessment of blood flow in tissue through the analysis of intensity fluctuations in diffusely backscattered coherent light. The non-invasive nature of DCS has enabled several clinical application areas for deep tissue blood flow measurements, including neuromonitoring, cancer imaging, and exercise physiology. While promising, in measurement configurations targeting deep tissue hemodynamics, standard DCS implementations suffer from insufficient signal-to-noise ratio (SNR), depth sensitivity, and sampling rate, limiting their utility. In this work, we present an enhanced DCS method called pathlength-selective, interferometric DCS (PaLS-iDCS), which uses pathlength-specific coherent gain to improve both the sensitivity to deep tissue hemodynamics and measurement SNR. Through interferometric detection, PaLS-iDCS can provide time-of-flight (ToF) specific blood flow information without the use of expensive time-tagging electronics and low-jitter detectors. The technique is compared to time-domain DCS (TD-DCS), another enhanced DCS method able to resolve photon ToF in tissue, through Monte Carlo simulation, phantom experiments, and human subject measurements. PaLS-iDCS consistently demonstrates improvements in SNR (>2x) for similar measurement conditions (same photon ToF), and the SNR improvements allow for measurements at extended photon ToFs, which have increased sensitivity to deep tissue hemodynamics (∼50% increase). Further, like TD-DCS, PaLS-iDCS allows direct estimation of tissue optical properties from the sampled ToF distribution. This method offers a relatively straightforward way to allow DCS systems to make robust measurements of blood flow with greatly enhanced sensitivity to deep tissue hemodynamics without the need for time-resolved detection, enabling further applications of this non-invasive technology.

## 1. Introduction

Although representing only ∼2% of the total body weight, the brain accounts for approximately 15%-20% of the cardiac output at rest [1]. Following acute brain injury, disruption in the supply of oxygen and glucose to the brain can have grave consequences and contribute to secondary brain injury, worse outcomes, and increased morbidity and mortality [2–4]. Current clinical methods used to assess cerebral blood flow (CBF) include magnetic resonance imaging (MRI) [5], computed tomography (CT) [6,7], and transcranial doppler ultrasound (TCD) [8,9], though none can provide continuous measurements of CBF. As an alternative, diffuse correlation spectroscopy (DCS), a diffuse optical technique used to estimate microvascular perfusion [10], allows for a non-invasive, continuous estimate of blood flow in the brain. DCS has been validated against several gold-standard perfusion monitoring techniques including arterial spin labeled MRI [11,12], Xenon-CT [13], positron emission tomography [14], and tracer bolus tracking [15], and has been used extensively in research to assess CBF in multiple clinical scenarios, including major cardiac surgeries [16–18] and long-term monitoring of acute brain injury in the neuro ICU [13,19,20]. DCS has also found use in other translational research areas including breast cancer imaging [21] and exercise physiology [22,23]. Traditionally, DCS has been performed using continuous-wave (CW) illumination, which, due to the partial volume effect, intrinsically links the measurement source-detector separation (SDS), cerebral sensitivity, and signal-to-noise ratio (SNR) of the measurement [24]. As a consequence, for source-detector separations sufficiently sensitive to the cerebral signal, the SNR of the measurement is limited, requiring slow sampling of the cerebral blood flow signal, and ultimately limiting the usefulness of the technique [25]. To address these limitations, several improvements to the basic DCS technique have been developed. To address insufficient SNR in CW-DCS, groups have developed DCS techniques based on longer wavelength light (i.e. 1064 nm) [26], massively parallelized signal detection [27–29], and the use of interferometry (iDCS) [30–33]. These methods have allowed for major improvements in measurement SNR at extended source-detector separations, though still maintain the link between source-detector separation, measurement SNR, and cerebral sensitivity. To decouple this link, methods based on time-of-flight (ToF) discriminative detection have also been developed and include time-domain DCS (TD-DCS) [34,35], interferometric near-infrared spectroscopy (iNIRS) [36], and coherence-gated DCS [37–39], enabling measurements taken at short source-detector separations that take advantage of the greater absolute number of photons that carry information about the cerebral hemodynamics signal as compared to long source-detector separations [40]. In this work, we synergistically combine several advances to DCS technology and introduce an improved coherence-gated DCS method termed pathlength-selective interferometric DCS (PaLS-iDCS). PaLS-iDCS uses long-wavelength illumination (1064 nm) and interferometric detection for both simultaneous time-of-flight selectivity and coherent gain, greatly improving measurement performance toward robust monitoring of cerebral blood flow beyond. Further, we demonstrate that by sweeping the coherence gate across the re-emitted light pulse we can also estimate sample optical properties without the need for a time-resolved detector.

## 2. Methods and Materials

### 2.1 Diffuse correlation spectroscopy

Diffuse correlation spectroscopy allows for the non-invasive estimation of blood flow in tissue through the analysis of fluctuations in coherent, diffusely back-scattered light [10]. To estimate blood flow, the temporal intensity autocorrelation function, *g*_2_(*τ*), of the fluctuating intensity signal is calculated and fit for an index of blood flow (BF_i_) for the tissue being measured [41]. The decay of the autocorrelation function of *g*_2_(*τ*) is related to the decay of the electric field, temporal autocorrelation function, *g*_1_(*τ*), by the Siegert relationship [42], given in Equation 1,

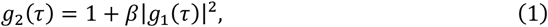

where *β* is the coherence parameter, which depends on the coherence length of the laser, the geometry of the measurement, and the degree of environmental light contamination. The electric field autocorrelation function is directly related to the blood flow in tissue due to the phase difference introduced by the dynamic scattering events. The electric field autocorrelation function for a single pathlength, s, at a time lag, *τ*, in a scattering medium is given in Equation 2 [41],

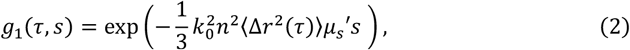

where k_0_ is the wavenumber, n is the index of refraction, ⟨Δ*r*^2^(*τ*)⟩ is the mean-squared displacement of the dynamic scattering particles, and 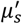 is the reduced scattering coefficient. For most experimental conditions, the measured mean-squared displacement is modeled as a diffusive process and takes the form ⟨Δ*r*^2^(*τ*)⟩ = 6*D*_*b*_*τ*, where D_b_ is the apparent diffusion coefficient. The BF_i_ is then estimated as the product of the apparent diffusion coefficient and the probability of scattering from a moving scatterer *α*, i.e. *BF*_*i*_ = *αD*_*b*_. For DCS measurements a distribution of pathlengths, P(s), which depends on tissue optical absorption and scattering as described by the established governing equations of light propagation in the diffuse regime [43], is collected, and the measured autocorrelation function, *g*_1_(*τ*) or *g*_2_(*τ*), reflects the weighting of the individual pathlength autocorrelation functions, *g*_1_(*τ, s*), given in Equation 3 for a continuous-wave (CW) DCS measurement made with a long coherence length laser,

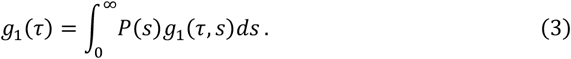

As mentioned in the introduction and demonstrated by the form of Equation 3, for measurements made in the CW configuration, hemodynamic information carried by different pathlengths is mixed. To reduce the influence of less cerebrally sensitive short pathlengths, extended source-detector separations (>∼30mm) are required for the influence of the cerebral signal to outweigh the influence of the extracerebral signal, and this results in a greatly attenuated measured light intensity [40,44]. Further, while continuous-wave illumination is the simplest implementation of near-infrared spectroscopic techniques, though some approaches are possible [45], information about tissue optical properties is difficult to extract. Without measured optical properties, the interpretability of the absolute blood flow index fit from the measured autocorrelations is then reduced.

### 2.2 Time-domain, diffuse correlation spectroscopy (TD-DCS)

To overcome the trade-off in cerebral sensitivity and light intensity (SNR) and to provide optical properties for greater interpretability of the extracted hemodynamic signals, DCS methods able to discriminate between photons of different pathlengths were developed. These have been based either on time-correlated single photon counting (TCSPC) [34], interferometric detection with a frequency swept source (iNIRS) [36], or low coherence interferometric detection [37,38]. Time domain diffuse correlation spectroscopy (TD-DCS) involves the use of sub-nanosecond pulses of light (100’s of ps), low jitter time-resolved single-photon detectors, and time-measurement electronics with ps-resolution. Detected photons are binned by their ToF into the temporal point spread function (TPSF), which allows for both the separation of “early photons,” which travel primarily in superficial tissue, from “late photons,” which have longer pathlengths and are more likely to carry information about cerebral perfusion, as well as permit the fitting of the shape of the TPSF for the absorption and scattering properties of the tissue, similar to time domain near-infrared spectroscopy [46]. ToF-specific selectivity allows for the use of shorter source-detector separations, which allows for an improved signal-to-noise ratio as mentioned in the introduction. For TD-DCS, the measured intensity autocorrelation function, given in Equation 4 [47], is expressed as,

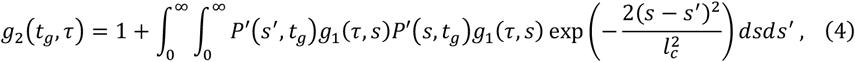

where t_g_ is the sample time of the measurement, *ν* is the speed of light in the medium, *l*_*c*_ is the coherence length of the laser, which for a transform-limited gaussian pulse is related to the pulse duration as 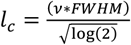 (derivation provided in supplemental section S1), and *P*^*′*^(*s, t*_*g*_) is the effective pathlength distribution, defined as 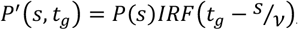, which is broadened by the instrument response function (IRF) and in practice is further broadened by the choice of photon selection profile (i.e. “gate”). The IRF of the system represents the temporal uncertainty of the arrival time of a particular photon that is due to the width of the input laser pulse and the uncertainty of the timing of photon detection, including detector non-idealities as well as timing jitter introduced by the detector and time tagging electronics. As has been shown previously [48], detector non-idealities, like the diffusion tail of many silicon SPAD detectors, can have negative consequences on the sensitivity of the measurement to blood flow in deeper tissue, as photons closer to the peak of the TPSF are erroneously registered as having occurred later in the total time of flight distribution. Further, due to the lopsided relative photon flux between the early time gates and the late time gates of interest (∼100x more early photons), if too short a source-detector separation or too low of a repetition rate of the laser is used at a given average input optical power, the probability of photon detection for each laser pulse will cross into a regime which is no longer governed by single photon statistics. In this case, detector hold-off time artifacts and the pile-up effect further reduce the photons collected at the later time gates, reducing SNR, and distorting the shape of the TPSF, which introduces errors in the estimation of optical properties. Finally, the need for low jitter detectors and fast detection electronics with resolution in the 10’s of picoseconds makes TD-DCS a more expensive technique than conventional diffuse correlation spectroscopy. To address these shortcomings, we have developed a coherence-gated method that achieves the same goals as TD-DCS while removing the need for time-resolved acquisition and improving measurement SNR.

### 2.3 Pathlength-selective, interferometric diffuse correlation spectroscopy (PaLS-iDCS)

To enable ToF-selective measurements without the use of fast timing electronics, we combine the previously developed interferometric diffuse correlation spectroscopy (iDCS) approach with the principles of TD-DCS. iDCS is characterized by the use of an interferometer which allows for the weak, diffusely backscattered light from tissue to be combined and amplified by a reference arm derived directly from the laser source [30]. This approach has the benefit of intrinsically improving the SNR of the measurement, as well as removing the need for single photon detectors through the increased magnitude of the measured speckle fluctuations. In Equation 5, we begin by expressing the general form of the PaLS-iDCS autocorrelation function, given as,

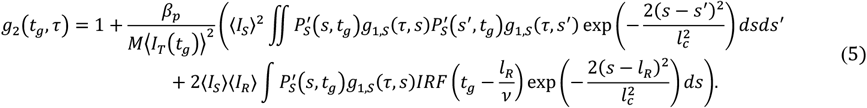

where *M* is the number of detected spatial modes, *β*_*p*_ is the polarization-dependent coherence factor, computed as 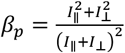, where *I*_∥_ and *I*_⊥_ represent the intensity carried by two orthogonal polarization components, ⟨*I*_*T*_(*t*_*g*_)⟩ is the average total intensity that falls within the selected time gate centered at *t*_*g*_, *l*_*R*_ is the length of the reference arm pathlength, the first integral term is the conventional TD-DCS autocovariance function (i.e. the integral expression in Equation 4 without the offset of 1) scaled by the square of the average total sample arm intensity, ⟨*I*_*S*_⟩, and the second integral term is the new PaLS-iDCS autocovariance term scaled by the product of the total sample intensity and the total reference intensity, ⟨*I*_*R*_⟩.

Under conditions in which the second term in Eq. 5 dominates, we can drop the requirement for time-resolved detection. Without fast time-tagging electronics, the information provided by the measurement time, *t*_*g*_, is lost, reducing the measurement from a pulsed TD-DCS system to a strictly CW one, though still reflecting a lower coherence length source. For PaLS-iDCS, due to the form of the autocorrelation function of the interference term, the pathlength-specific autocorrelation function and information about deep tissue blood flow can be encoded even without fast-timing information, reducing the complexity of the acquisition electronics to that of a typical DCS measurement, (in fact potentially simpler, since the reference boosted interference signal no longer necessitates photon counting detection [30,31]), while maintaining the benefits of the sensitivity to deep flow of the time domain approach. This benefit also extends to the estimation of optical properties. By evaluating the expression of the form of the autocorrelation at zero-time lag, the optical properties of the tissue can be estimated from the overall coherence parameter, *β*(*t*_*g*_), of the autocorrelation function for the PaLS-iDCS measurement measured at multiple gate times (*t*_*g*_) using the model given in Equation 6,

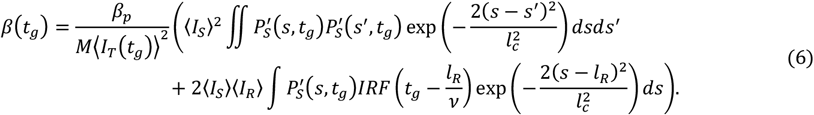

Full derivations of Equations 5 and 6 are found in supplemental section S.2.

### 2.4 Description of the multi-layer Monte Carlo simulations

To evaluate the theoretical performance of PaLS-iDCS and compare it to the theoretical performance of TD-DCS, multi-layer Monte Carlo simulations were performed. A three-layer model consisting of the scalp, skull, and brain is adopted for these simulations. In previous work, three-layer models omitting the CSF have been found to recapitulate both simulated and real measurements with only marginal observed differences to a four-layer model where CSF is included [36,49] and additionally provide more accurate representations of tissue than do two layer models which lump scalp and skull due to the high difference in perfusion between scalp and skull [36]. For both techniques, to allow for comparison to both phantom and human subject experiments at a count rate of the TD-DCS measurement that respects single photon statistics, a source-detector separation of 2 cm was used. The IRF of the system is assumed to be determined only by the width of the pulsed laser, neglecting the jitter introduced by the optical detector and related electronics. This situation represents the most optimistic situation for TD-DCS, with realistic values of jitter only leading to the degradation of the performance expected from the reported values. As mentioned previously, PaLS-iDCS will be unaffected by detection jitter. Different pulse durations were tested between 100 ps and 500 ps to determine optimal values of pulse duration for the measurement geometry. We also explored gate durations of 2/3, 5/3, and 10/3 of the pulse duration, as was done in a previous TD-DCS work [50]. For the PaLS-iDCS simulations, the gate duration is the entire TPSF, demonstrating the pathlength selectivity without the need for fast timing information. The simulated laser pulse is assumed to be transform limited, and the coherence length used for each simulation is computed as a function of the pulse duration. The reference arm was set to have a total intensity that was a factor of 10^4^ greater than the intensity contained within the entire sample TPSF to allow for the condition in which the contribution of the sample arm to the shape of the measured autocorrelation function is negligible. A range of gate center times was explored from 500 ps before the peak of the TPSF to 2100 ps after the peak of the TPSF. Simulated noise based on the previously described correlation noise model [51] was added to the autocorrelation functions to allow for the comparison of the noise properties of the techniques. A summary of the relevant parameters explored in the Monte Carlo simulations can be found in Table 1 and Figure 1a.

**Table 1.**
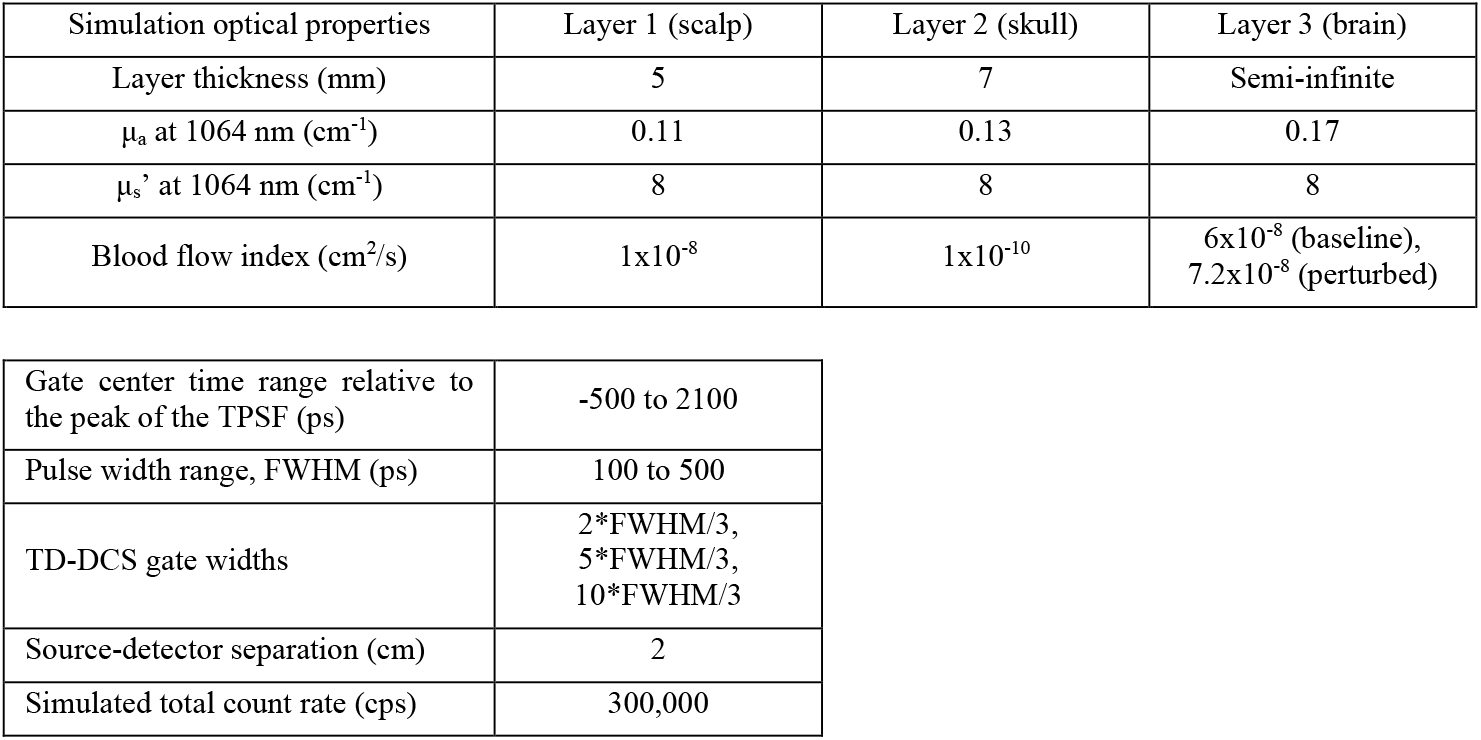
Monte Carlo simulation parameters.

| Simulation optical properties | Layer 1 (scalp) | Layer 2 (skull) | Layer 3 (brain) |
| --- | --- | --- | --- |
| Layer thickness (mm) | 5 | 7 | Semi-infinite |
| $\mu_a$ at 1064 nm ($\text{cm}^{-1}$) | 0.11 | 0.13 | 0.17 |
| $\mu_s'$ at 1064 nm ($\text{cm}^{-1}$) | 8 | 8 | 8 |
| Blood flow index ($\text{cm}^2/\text{s}$) | $1 \times 10^{-8}$ | $1 \times 10^{-10}$ | $6 \times 10^{-8}$ (baseline),<br>$7.2 \times 10^{-8}$ (perturbed) |

|  |  |
| --- | --- |
| Gate center time range relative to the peak of the TPSF (ps) | -500 to 2100 |
| Pulse width range, FWHM (ps) | 100 to 500 |
| TD-DCS gate widths | 2*FWHM/3,<br>5*FWHM/3,<br>10*FWHM/3 |
| Source-detector separation (cm) | 2 |
| Simulated total count rate (cps) | 300,000 |

**Figure 1:**
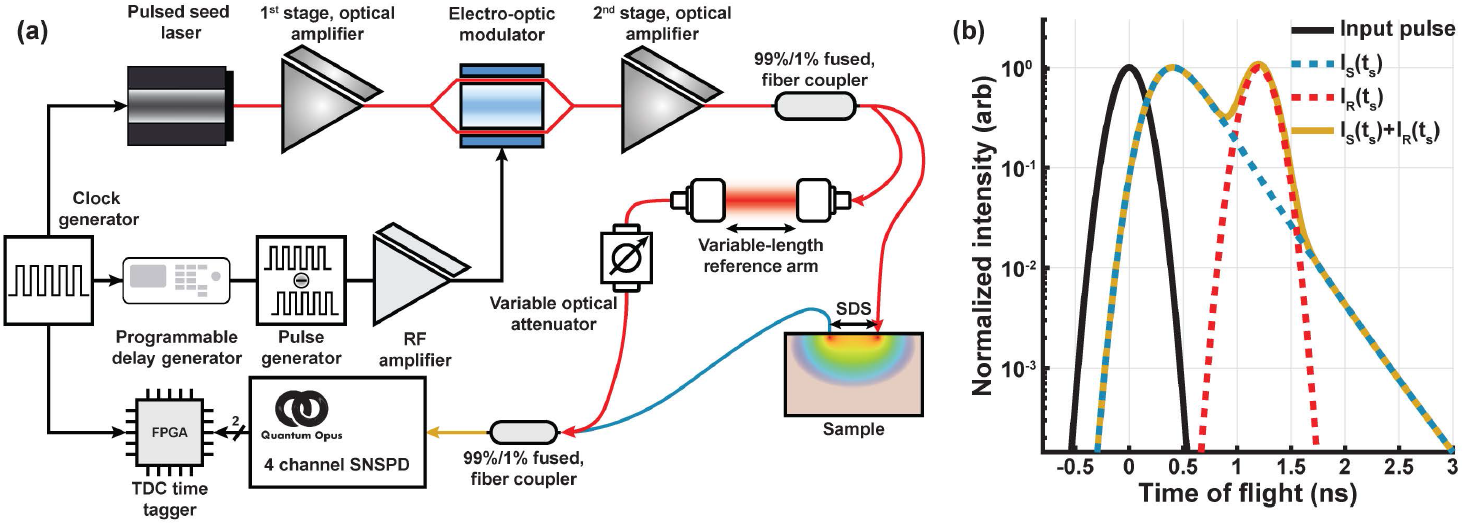
(a) Depiction of the PaLS-iDCS experimental setup. The output of the previously described custom pulsed laser source is split into sample and reference arms and the diffusely reflected light is collected from the sample and recombined with the delayed and attenuated reference arm. In (b) the time-of-flight distribution of the input pulse, the return from the sample (I_S_(t_g_)), the reference arm(I_R_(t_g_)), and the combined distribution of the sample and reference arms. The position of the reference arm within the sample time-of-flight distribution determines the time-of-flight selection.

To compare the simulated performance between TD-DCS and PaLS-iDCS, several metrics were computed, including: cerebral sensitivity, defined as 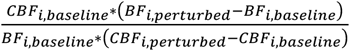 where the CBF terms are based on the ground truth BF_i_ in the cerebral layer and the BF_i_ terms are the model fits; coefficient of variation (CoV) of the fit BF_i_, defined as the ratio of the standard deviation of the BF_i_ fit to the mean value of the BF_i_ fit; and contrast-to-noise ratio (CNR) for the cerebral signal, defined as the sensitivity divided by the CoV. These factors are computed for each gate center time, pulse width, and gate width for both methods. A single perturbation of a 20% increase in cerebral blood flow is utilized to assess sensitivity, with sensitivity derived from single perturbations being shown to provide roughly equivalent estimates of sensitivity across a range of perturbation amplitudes [44]. For reference, to compare to the state-of-the-art for DCS with CW illumination, a simulation of iDCS performance at a range of source-detector separations between 5 mm and 40 mm in increments of 5 mm was performed to evaluate the performance of the time-resolved techniques to the performance of a CW-iDCS system. Autocorrelation functions in this work were fit using the full model for the TD-DCS and PaLS-iDCS autocorrelation functions expressed in Equations 4 and 5, respectively. To limit complexity, as is common in the field, the simulated data are fit assuming a semi-infinite medium (i.e. homogeneous, no layered structure) with optical properties μ_a_ = 0.14 cm^-1^ and μ_s_’ = 8.0 cm^-1^.

### 2.5 Description of the TD-DCS and PaLS-iDCS instrumentation

Shown in Figure 1, the measurement system consists of a custom 75 MHz amplified and shaped pulsed 1064 nm laser source described previously [50], a Mach-Zehnder interferometer with a variable length reference arm, and superconducting nanowire single photon detectors (SNSPDs). In this work, we utilize detectors with optimal timing characteristics, i.e. gaussian single-photon response with <100 ps (FWHM) timing resolution, to demonstrate and validate the PaLS-iDCS technique. The amplified, pulsed-shaped source light is split into sample and reference arms using a 99%/1% fused fiber coupler (PN1064R1A2, Thorlabs). The 99% arm is coupled into a 400 µm-core fiber (FT400EMT, Thorlabs) to deliver the maximal permissible exposure limit of 100 mW in a 3.5 mm aperture at 1064 nm as set by the ANSI standard (Z136.1). The 1% arm is coupled to a fiber collimator (PAF2A-15B, Thorlabs) at one end of the variable length reference arm inside an optical cage. Another identical fiber coupler is placed opposing the first, and this arrangement forms the variable length reference arm. The variable length reference arm has a translation range of ∼50 cm (1.66 ns), allowing for the reference pulse to be swept across most of the TPSF. In this proof-of-concept implementation the reference arm adjustment is manual, though we note that the processes can be substantially accelerated in the future through automation using either motorization or fiber optic switching. A single-mode fiber collects the reference arm light via the second fiber collimator and was fusion-spliced to the 1% arm of a 99%/1% fused fiber coupler for recombination with the sample return light. Co-located single mode detection fibers (780HP) were placed at a source-detector separation of 2 cm on the sample. The TD-DCS detector fiber was coupled directly into the SNSPD detector (Opus One, Quantum Opus), and the PaLS-iDCS detector fiber was fusion spliced to the 99% arm of the aforementioned 99%/1% fused fiber coupler (TN1064R1A2A, Thorlabs). The combined output of the coupler was routed to a second SNSPD channel. Electrical pulses corresponding to photodetection events were provided by the SNSPD system output and directly fed to two independent inputs of an FPGA-based time-tagger unit (Time Tagger Ultra, Swabian Instruments) together with a copy of the laser trigger signal for TCSPC reconstruction. Acquired data were stored in the instrument computer for post-processing.

### 2.6 Description of phantom experiments

Liquid phantoms were made mixing water and 20% intralipid (Fresenius Kabi) to reach a reduced scattering coefficient (μ_s_’) of ∼6.5 cm^-1^. The absorption coefficient (μ_a_) of the phantom at 1064 nm was ∼0.12 cm^-1^, driven by the absorption coefficient of water. Shaped pulses of FWHM durations of 100 ps, 200 ps, 300 ps, 400 ps, and 500 ps were compared for signal-to-noise ratio across measurements. PaLS-iDCS measurements were taken moving the reference arm from an initial position of ∼300 ps before the peak of the TPSF to ∼1000 ps after the peak of the TPSF in steps of ∼167 ps (5 cm in air). Autocorrelation functions for both techniques were calculated at a sampling rate of 10 Hz. Comparisons of signal-to-noise ratio of the fit BF_i_ between concurrently measured TD-DCS signals, at the same gate after the peak of the TPSF, were made. In addition, optical property estimates of the phantom were derived from both TD-DCS and PaLS-iDCS measurements. For TD-DCS, the TPSF was fit for both μ_a_ and μ_s_’ using the theoretical model for the pathlength distribution of diffusely backscattered light at a given source-detector separation in the semi-infinite reflectance geometry and the IRF of the system. For PaLS-iDCS, the optical properties of the phantom are fit from the reference arm pathlength sweep using the relationship described in Equation 6 between the value of the coherence parameter of the normalized autocorrelation function, the pathlength distribution, reference arm position, system IRF, and the coherence properties of the source. For both techniques, data from between 200 ps before the peak of the TPSF to 1.1 ns after the peak of the TPSF was used to fit for the optical properties.

### 2.7 PaLS-iDCS demonstration in human subjects

We experimentally validated the improvements in signal-to-noise ratio using PaLS-iDCS on five healthy volunteers. This study was reviewed and approved by the Mass General Brigham Institutional Review Board (#2019P003074). All participants gave written informed consent prior to the measurements. All methods were performed in accordance with the relevant guidelines and regulations. To compare the performance of TD-DCS and PaLS-iDCS directly, co-located detector fibers were placed at a source-detector separation of 2 cm, and a short source-detector separation channel at 5 mm was also added (Perkin Elmer, SPCM-AQR-14FC). A shaped pulse with FWHM of 300 ps was used for the human subject experiments. The sample arm photon count rate was ∼250 kcps for each subject at the 2 cm source-detector separation, allowing for a high reference-to-sample arm count rate ratio that still obeyed single photon statistics. To better visualize pulsatile hemodynamic oscillations, autocorrelation functions were calculated at an increased rate of 20 Hz. A baseline resting measurement of 60 s was taken for different lengths of the PaLS-iDCS reference arm to evaluate the *in vivo* performance of the technique at different time gates. The reference arm was translated to reach gates that were 200 ps before the peak of the TPSF to ∼1 ns after the peak of the TPSF in steps of 133 ps. BF_i_ values sampled at 20 Hz, allowing for the visualization of pulsatile blood flow, as well as the coefficient of variation of the fit BF_i_, are compared between TD-DCS and PaLS-iDCS for each of the selected gates. The pathlength sweep was taken over 30 minutes in each subject. Additionally, to evaluate increases in cerebral sensitivity *in vivo*, a pressure modulation procedure was performed to selectively reduce the blood flow in the skin and assess changes in the measured blood flow index at different time gates and the short separation channel [52]. To accomplish this, an elastic band was placed just below the probe and pulled tight against the forehead. The procedure consisted of a 60 s baseline, three repeated sets of 60 s of pressure and 60 s of recovery, and an additional 60 s of recovery. The relative decrease in the blood flow index (rBF_i_) at the 5 mm short separation was used as reference for the scalp blood flow changes. By comparing the rBF_i_ for later photon time gates to the reference signal, an estimate of the scalp sensitivity of the measurement can be made, and by extension, from a previously observed relationship, the brain sensitivity of the measurement. Comparisons of both the pulsatile feature changes as well as slow changes (signals down sampled to 1 Hz) are performed between the short separation channel and late time gate.

## 3. Results

### 3.1 Comparison of the simulated performance between TD-DCS and PaLS-iDCS

In Figure 2, the comparison of simulated sensitivity (2.c and 2.f), coefficient of variation (2.d and 2.g), and contrast-to-noise ratio (CNR) (2.e and 2.h) between TD-DCS and PaLS-iDCS is shown as a function of gate position relative to the peak of the temporal point spread function (TPSF). To compare more cleanly across different pulse durations, as opposed to comparing parameters as a function of time relative to the peak of the TPSF, the CNR is compared across pulse durations as a function of the cerebral sensitivity. The optimal CNR operating condition at each level of cerebral sensitivity for both techniques is extracted and overlayed with the results of the CW-iDCS simulations obtained by changing source-detector separation (Figure 2.b). CW-iDCS cerebral sensitivity (x-axis) is directly proportional to source-detector separation, with the 5 mm SDS exhibiting almost no sensitivity to changes in the cerebral layer (leftmost purple diamond) and the 40 mm SDS exhibiting ∼45% sensitivity to the cerebral layer (rightmost purple diamond), with intermediate distances, 10 mm through 35 mm, ordered in increasing sensitivity. PaLS-iDCS provides a peak CNR operating point that is higher than both TD-DCS and CW-iDCS and exhibits a higher sensitivity to the cerebral signal at the optimal operating condition. As was observed previously [53], the CW technique provides a higher peak CNR than TD-DCS, though TD-DCS still provides higher CNR at operating points with increased sensitivity to the cerebral signal. The expanded results shown for PaLS-iDCS (Figure 2.c, 2.d, 2.e) and TD-DCS (Figure 2.f, 2.g, 2.h) demonstrate the influence the selection of gate position and pulse duration have on each of the simulation-derived parameters. The selection of optimal pulse duration, shown in Figure 2.b, is seen to alternate between two pulse durations in a relatively short range of the cerebral sensitivity value. This alternation is due to the similarity in shape of the CNR curves for each pulse duration as a function of time-of-flight (Figure 2.e and 2.h) when the curves are shifted in time-of-flight to maximize the overlap in the corresponding sensitivity curves (Figure 2.c and 2.f), making the selection of each pulse duration effectively equivalent. In addition to gate position, for TD-DCS, the selection of gate width has an influence on the performance of the measurement. Figures 2.f, 2.g, and 2.h display results for a gate which is 5/3 the width of the full-width at half maximum (FWHM) of the pulse, which was the optimal gate choice of the three tested durations for each of the pulse durations (supplement Figure S3). From these results, for PaLS-iDCS at a 2 cm source-detector separation with optical properties similar to those in tissue, a pulse duration of 300 to 400 ps will optimize the tradeoffs between sensitivity and signal-to-noise ratio, allowing for improved measurement of cerebral hemodynamics.

**Figure 2:**
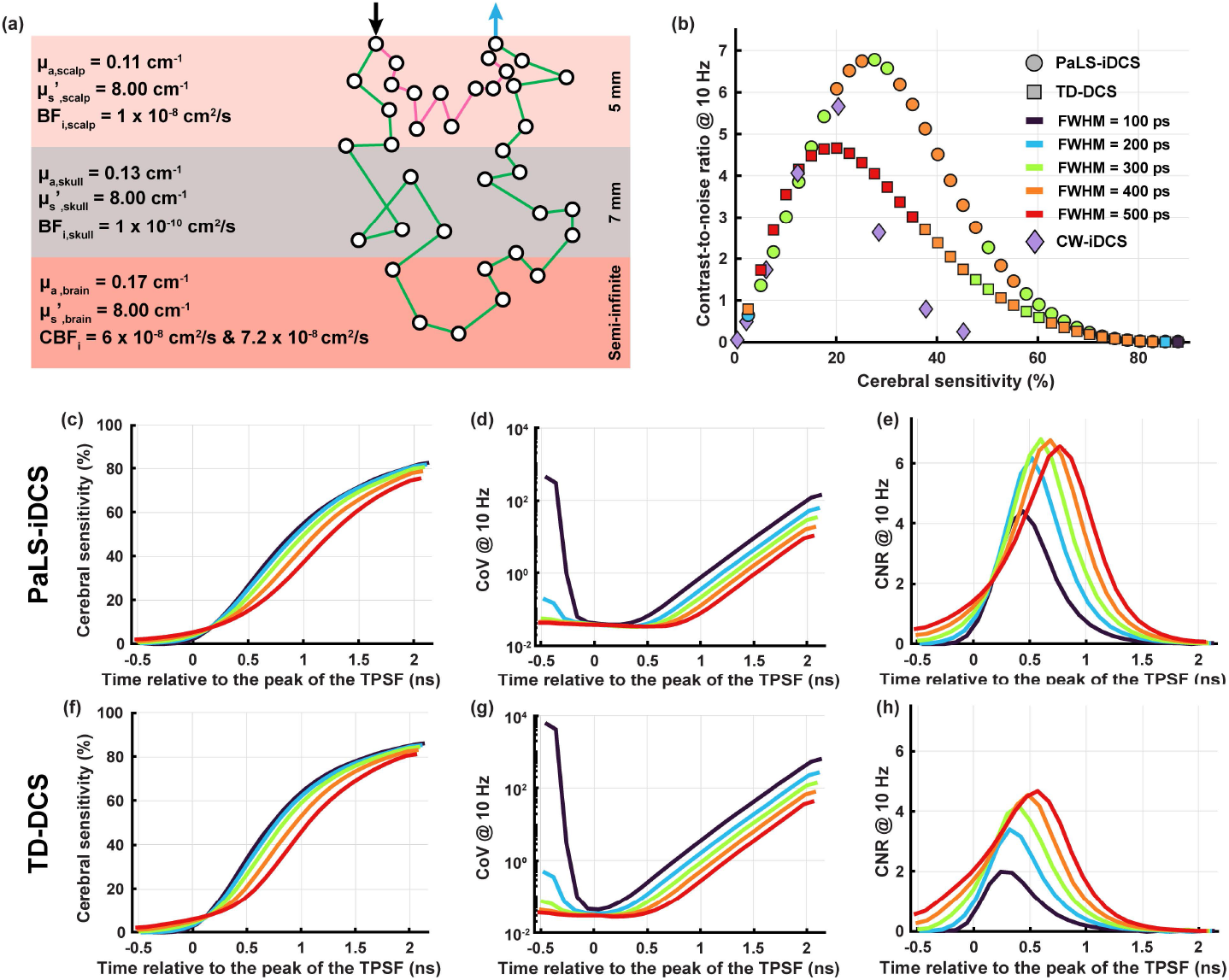
(a) Depiction of the tissue model used for the Monte Carlo simulations with labeled optical and perfusion properties. A short path (pink) and long path (green) through the tissue are shown to illustrate the improved sensitivity to deeper tissue that comes from selecting longer time-of-flight photon detections. The summarized results comparing CNR for the cerebral hemodynamic signal as a function of cerebral sensitivity are shown in (b). This comparison also includes CW-iDCS (diamonds). For CW-iDCS, the sensitivity is proportional and monotonically increasing with source-detector separation with the leftmost diamond representing the sensitivity and CNR at 5 mm and the rightmost diamond representing the sensitivity and CNR at 40 mm. PaLS-iDCS (circles) provides both the absolute highest CNR of the three compared techniques, as well as providing the highest CNR for measurements which exhibit >20% sensitivity to the cerebral signal. The pulse duration which achieved the CNR result for PaLS-iDCS and TD-DCS (squares) is encoded through the color of the marker. In (c-e) and (f-h) the results for PaLS-iDCS and TD-DCS, respectively, are shown as a function of the selected gate center time relative to the peak of the TPSF for the different pulse widths. As a function of gate center time, TD-DCS provides a slightly higher (∼5%) sensitivity for each pulse duration when compared to PaLS-iDCS (c vs. f). This small difference in sensitivity is outweighed by the reduction in coefficient of variation (CoV) at later times after the peak of the TPSF by PaLS-iDCS (d vs. g), which results in an overall improvement of CNR for each pulse duration at each gate center time after the peak of the TPSF (e vs. h).

### 3.2 Comparison of measurement performance between TD-DCS and PaLS-iDCS in phantoms

In Figures 3.b and 3.c, the comparison between the fit for BF_i_ at different times of flight is shown for TD-DCS and PaLS-iDCS, respectively. Consistent with the optimal gate duration found in simulation, the gate duration for both techniques is set equal to 5/3 the FWHM of the pulse. From the estimates of the diffusion coefficient calculated using the full autocorrelation model, while both techniques provide similar BF_i_ values when averaging across all gate center times and pulse durations (TD-DCS: (1.74±0.19)x10^−8^ cm^2^/s, PaLS-iDCS: (1.80±0.10)x10^−8^ cm^2^/s), the PaLS-iDCS technique provides a more consistent estimate of the diffusion coefficient across pulse durations and gate center times. This may reflect non-idealities that come with a real measurement system, including jitter in the photon detection timing introduced by the detector and the timing electronics as well as uncertainty in the mutual coherence function of the laser pulse at different ToF displacements (supplemental section S3), both of which can affect the estimate of the contributions of different pathlengths to the resultant autocorrelation. By utilizing the coherence-gated approach, the estimate of the diffusion coefficient is more consistent, even with the non-idealities present in the measurement system. In Figures 3.d and 3.e, the signal-to-noise ratio (SNR) of the plateau of the correlation function (*g*_2_(*τ*)) and the coefficient of variation (CoV) of the fit diffusion coefficient are compared between TD-DCS and PaLS-iDCS, respectively. For all gate times and pulse durations, PaLS-iDCS provides both higher SNR and reduced CoV as compared to TD-DCS, consistent with the observations seen in the simulations. The SNR observed in the measurements also maintains the relationship observed with respect to pulse duration, with PaLS-iDCS SNR peaking at a time within the interval of tested pulse durations, and the TD-DCS SNR continuing to increase with increasing pulse duration.

**Figure 3:**
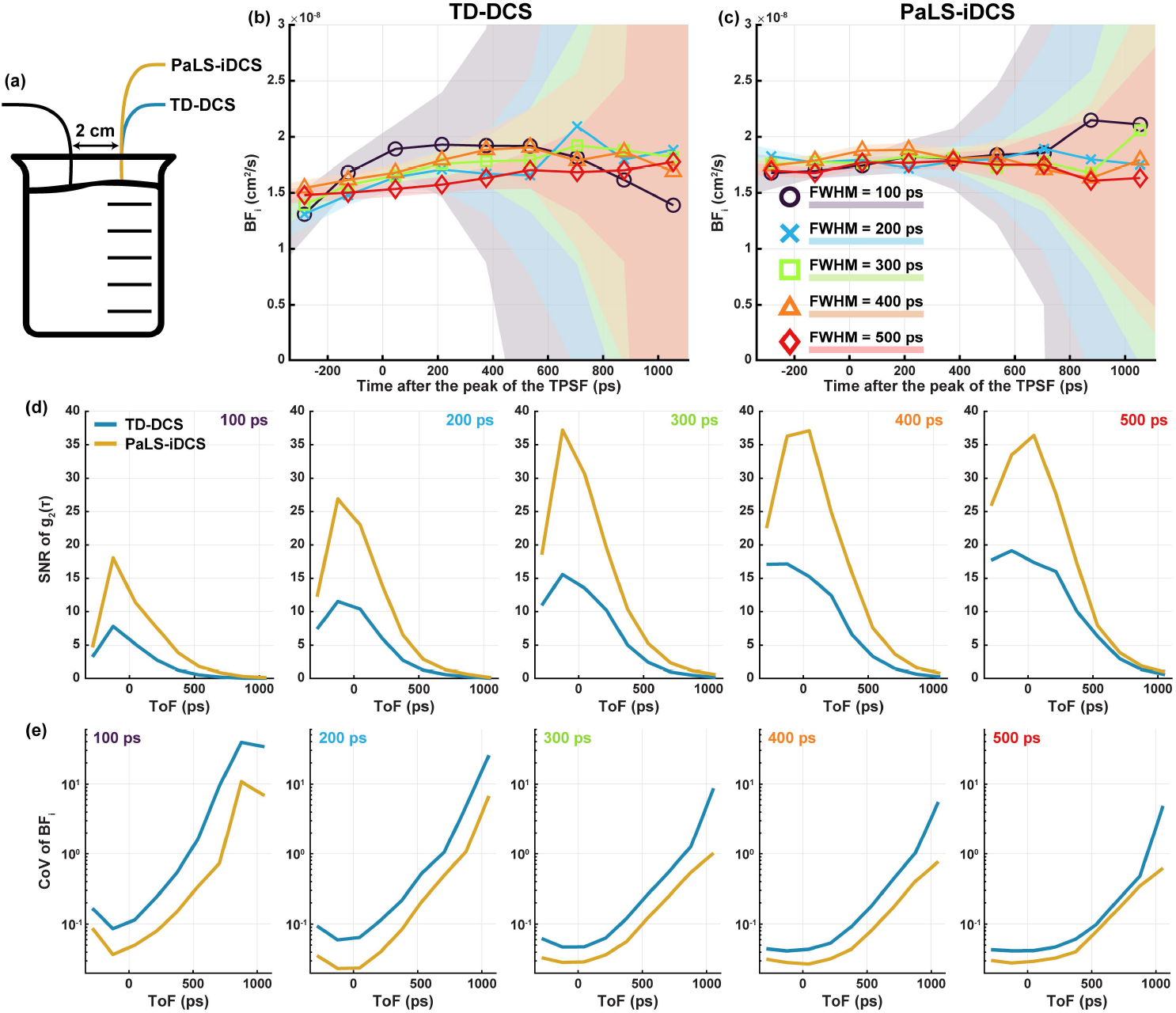
(a) Depiction of the intralipid phantom measurement with a source-detector distance of 2 cm. In (b) and (c), fit values of BF_i_ are shown for TD-DCS and PaLS-iDCS, respectively (shading indicates ±1 standard deviation from the mean). For comparisons within a pulse duration and across pulse durations, PaLS-iDCS can be seen to give more consistent results for the estimation of the BF_i_ of the phantom with a reduced variability at a given gate center time relative to TD-DCS, as shown by the narrower shaded bands. The SNR of the autocorrelation functions (d) and CoV of the BF_i_ (e) at a 10 Hz acquisition rate demonstrates the improvement in performance of the PaLS-iDCS (yellow lines) instrument over the TD-DCS (blue lines) instrument.

Optical properties derived from the phantom experiments at different pulse durations are found to be similar between both techniques and across pulse durations, as seen below in Figure 4, for both absorption (Figure 4.b) and scattering (Figure 4.d). The PaLS-iDCS derived optical properties are relatively noisier (the error bars represent ±1 standard deviation), which is likely related to the relatively sparse sampling of the measured beta values due to the manual adjustment mechanism used for reference arm length adjustment. The TPSF provided by the time-tagging electronics is much more densely sampled with a bin width of 10 ps (Fig 4.a, plotted at 50 ps spacing) as compared to the interferometric beta values, which are sampled every 167 ps (Fig 4.c). Though noisier in this demonstration, the ability of the PaLS-iDCS technique to assess optical properties from a reference arm pathlength sweep can still significantly improve the quantification of blood flow vs. a “classic” CW-DCS measurement by providing accurate optical properties to the fitting model without the use of time-correlated single photon counting (TCSPC) acquisition and potentially without needing photon counting detection given the amplified signal level at the photodetector.

**Figure 4:**
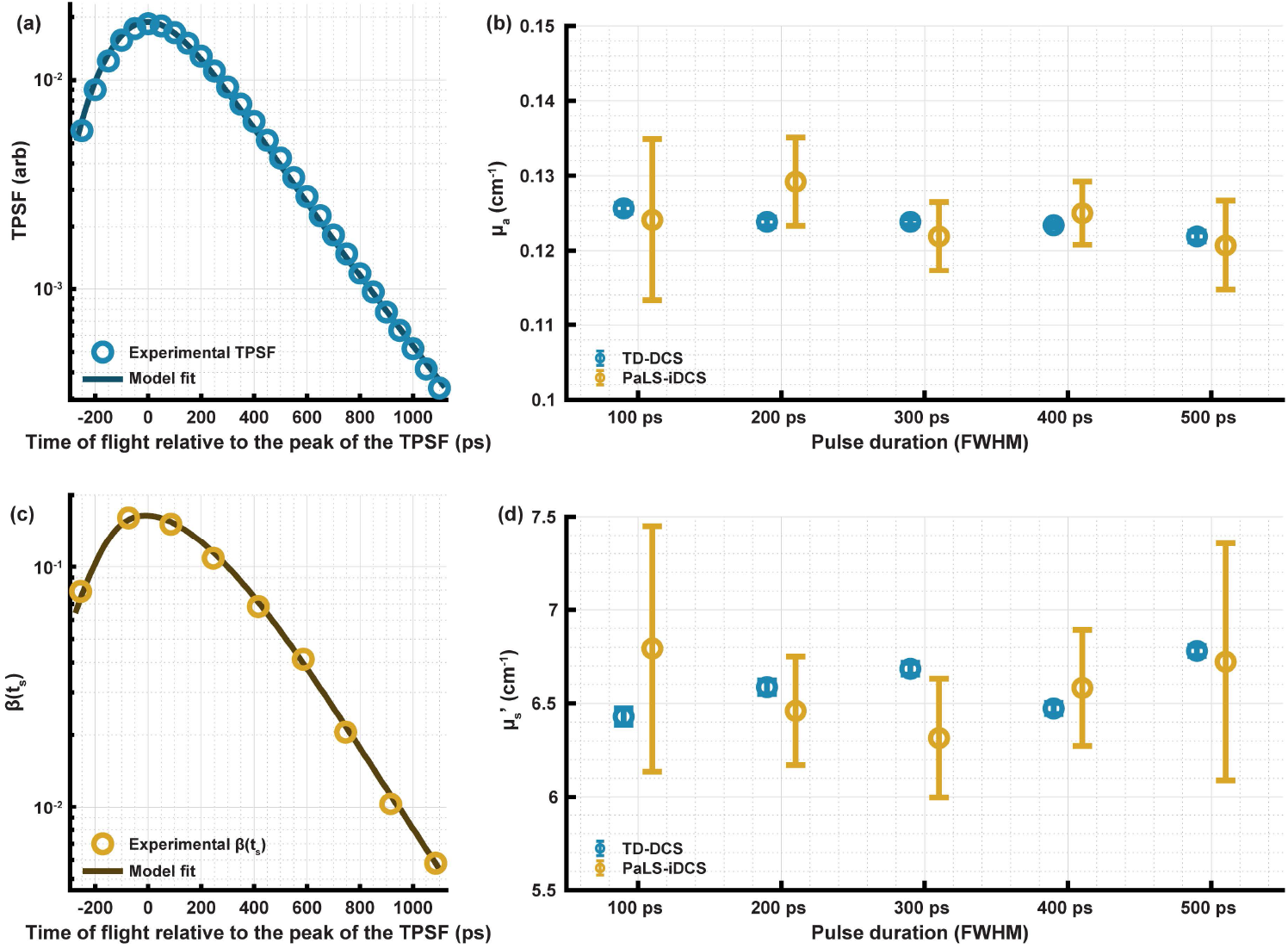
Examples of the data fit for the optical properties for TD-DCS (a) and PaLS-iDCS (c). TD-DCS fitting is related directly to the shape of the collected TPSF, analogous to time domain NIRS, while PaLS-iDCS is fit with the model expressed in Equation 6. Both the absorption (b) and scattering (d) properties fit from both techniques across pulse durations are generally in agreement. The PaLS-iDCS measurement of optical properties for both absorption and scattering are noisier than the measurements made by TD-DCS, though denser sampling of the reference arm sweep closer to that of the temporal sampling of the TPSF may improve the precision of the estimates.

### 3.3 Comparison of measurement performance between TD-DCS and PaLS-iDCS in healthy subject measurements

To further demonstrate the improved performance of PaLS-iDCS, we applied this technique in five healthy volunteers. By increasing the length of the reference arm, the information collected by the system increasingly reflects the longer pathlength light, which is more sensitive to the cerebral signal. In Figure 5.a, we demonstrate the improved SNR of the pulsatile signal as a function of time-of-flight selected at 133 ps increments in an example subject, with the gates adopted depicted in Figure 5.b. The left column shows BF_i_ time courses for 10 s of data acquired at rest; the right column shows the reconstructed pulsatile component averaged over the 60 s measurement duration. The time courses can be seen to become noisier with increasing time-of-flight. Though noisy, PaLS-iDCS provides recognizable cardiac pulsatility on a beat-by-beat basis (Fig 5.a) at an extended time gate of 800 ps after the peak of the TPSF at a 20 Hz sampling rate. Comparing the coefficient of variation of the estimated reconstructed pulsatile signal, with the interferometric technique, the same measurement CoV can be reached at a time of flight ∼300 ps later when compared to TD-DCS (Figure 5.c). Estimating from the simulations, this represents a roughly 80% increase in relative cerebral sensitivity. Additionally, from the full sweep of the reference arm, the optical properties measured by PaLS-iDCS provide similar values to the optical properties measured by TD-DCS (Figure 5.d). These performance improvements are also accompanied by the benefit of not requiring a time-of-flight sensitive detector.

**Figure 5:**
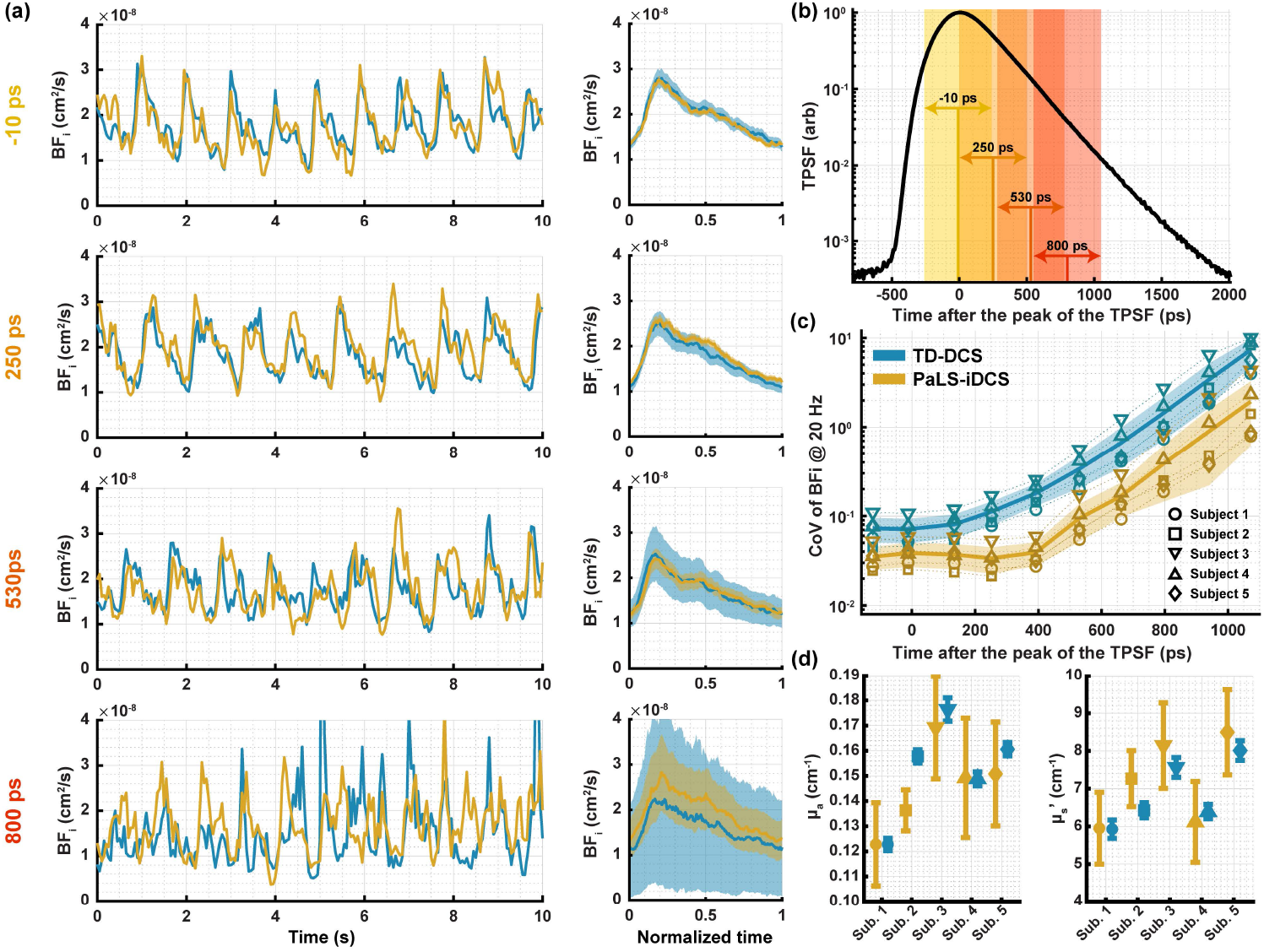
(a) Comparison of the time courses of BF_i_ for co-located TD-DCS (blue lines) and PaLS-iDCS (yellow lines) measurements for different photon selection gates. In addition to the time course comparison, the average pulsatile waveform (solid lines) as well as uncertainty (±1 std, shaded area) in the fit is shown for each of the photon selection gates. At each of the selected photon selection gates (shown in (b) with shaded areas and labeled with the gate center time corresponding to the waveforms shown in (a)), PaLS-iDCS provides a consistent improvement in the coefficient of variation of the fit of BF_i_ and allows for measurements at times of flight where the TD-DCS measurements begin to fail (i.e. 800 ps). (c) The distribution of the coefficient of variation at different points in the cardiac cycle for the BF_i_ fit at 20 Hz is approximately 2.5x to 4x lower for PaLS-iDCS than TD-DCS at each of the tested time of flight gates. (d) As in the phantom experiments, using the full pathlength sweep of the reference arm, PaLS-iDCS provides estimates of the optical properties of the tissue that are similar to those as estimated by TD-DCS.

In Figure 6 we present the results obtained in the pressure modulation experiment using a reference arm pathlength that selected a gate 750 ps after the peak of the TPSF. After identification of the pulsatile BF_i_, average BF_i_ within each pulse boundary was calculated, and the signals were resampled to a 1 Hz sampling rate to improve the SNR. Figure 6.a shows the collected TPSF in this condition with both the early (green) and late (red) gates shaded to indicate photon selection. We analyzed the difference between the blood flow response measured from the short separation and from photons arriving before the peak of the TPSF as compared to the extremely late gate time. Across all subjects, results from pressure modulations where the short separation BF_i_ was reduced by more than 50% were averaged together and time courses of the block averaged rBFi are shown in Figure 6.b, with the gray area representing the pressure modulation task. Figure 6.c summarizes the percent reduction of relative blood flow (rBF_i_) during the task. To address possible differences with each pressure modulation trial, the results are presented with respect to the reduction in short separation rBF_i_ that was seen in each trial. The unnormalized results can be found in supplemental section S6. The reduction at the late gate is ∼20% less than the reduction seen at the early gate, demonstrating the reduced sensitivity to superficial contamination and an increase in the sensitivity to the cerebral signal. Because these measurements are made at 2 cm, even at the peak of the TPSF there is still some sensitivity to the cerebral signal, though the ∼20% improvement at late gates is significant.

**Figure 6:**
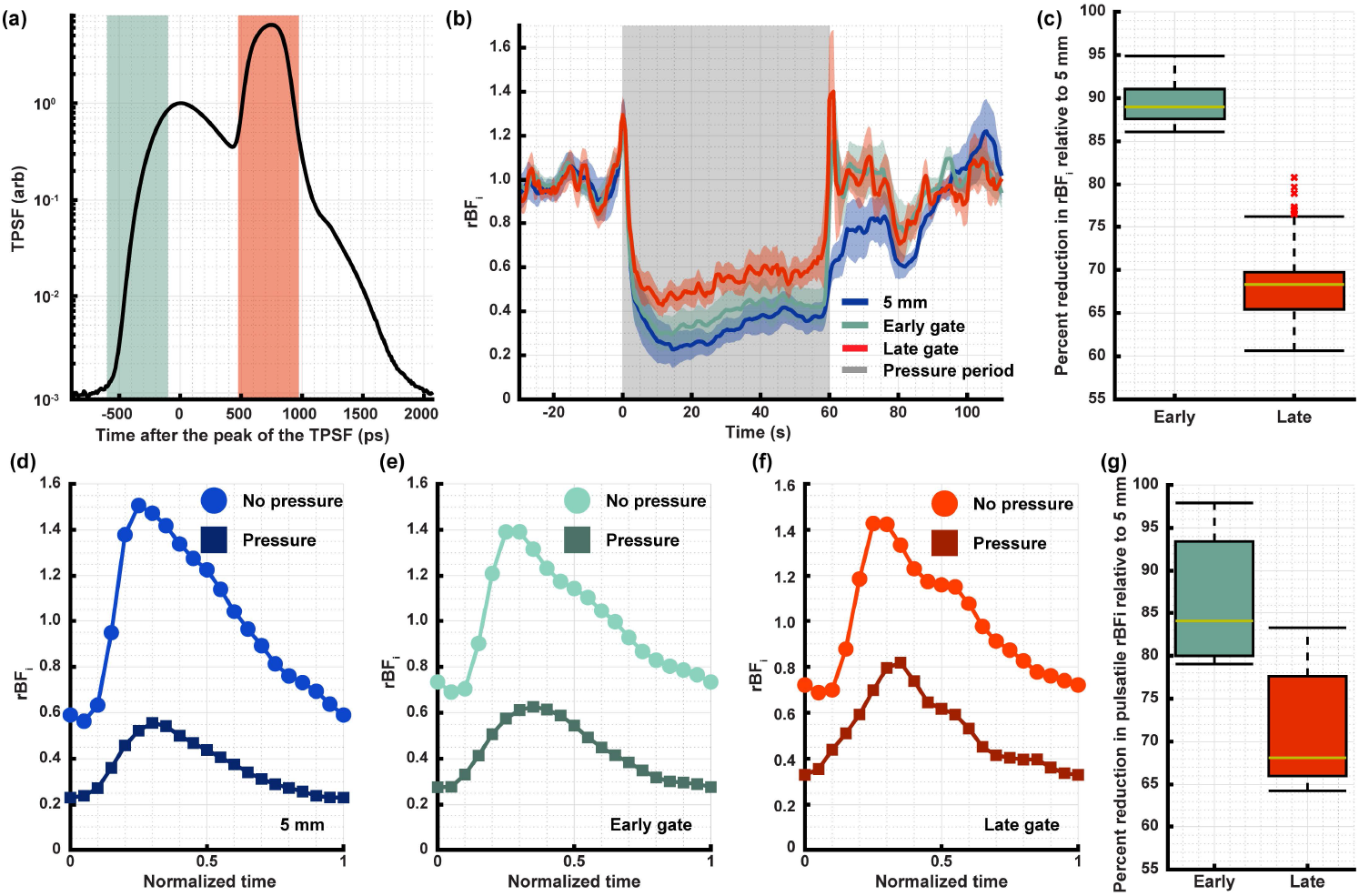
(a) The TPSF of the combined sample and reference arms is overlaid with the photon gate selection for the early (pre-peak, green shading) gate and the late (reference arm selected, red shading) gate. In (b) the block averaged change in relative BF_i_ is shown for the short separation (blue line), early (green line) and late (red line) gates with the standard error between trials shown as the shaded regions, and the distribution of the reduction in blood flow index for each gate is shown in (c), demonstrating the benefits of the later gate during the pressure modulation task (gray shading). Additionally, because the PaLS-iDCS instrument provides high SNR at the later gate times, in (d), (e), and (f) we show the average pulsatile waveform from the baseline period, labeled as no pressure (circles), and the average pulsatile waveform from the pressure modulation period (squares) to compare changes in pulse amplitude and pulse morphology for an example subject. The changes in pulse amplitude are quantified for all pulses in the pressure period across all subjects for all valid trials, and the distribution of the reduction in the amplitude is shown in (g).

Additionally, we can assess the difference in the reduction in pulsatile amplitude as a function of time-of-flight by averaging all detected heartbeats during each experimental period. As was the case for the average flow change, the later gate (Figure 6.f) exhibits a better-preserved pulsatile amplitude as compared to the early gate (Figure 6.e) or the short separation (Figure 6.d), indicating a lesser degree of superficial contamination of the pulsatile waveform shape, as summarized in Figure 6.g.

## 4. Discussion and conclusion

We have demonstrated a novel method for the measurement of optical properties and pathlength-selective flow using a pulsed laser and a coherence gate. While this is not the first DCS coherence-gating demonstration, previous techniques have either tested extremely short coherence length sources or comparatively longer coherence length sources, which are suboptimal in terms of signal-to-noise ratio and sensitivity to the cerebral signal. The method and tunable laser source described in this work have the potential to enable optimal measurements of cerebral blood flow under real world scenarios when measuring healthy subjects and patients. For the range of pulse durations tested and corresponding widths of the coherence gates, in the measurements shown here and in the supplementary information, the enhanced DCS system is capable of producing the optimal laser pulse duration of 300 to 400 ps. In comparison, in Zhao et al. [37], a relatively long coherence length source was created through the MHz modulation of a long coherence CW source. The generated time-of-flight filter, a feature that allows for coherence gating in a similar fashion to the placement of the reference arm pulse, had a FWHM that was between ∼1.2 and ∼1.6 ns depending on the time-of-flight of the center of filter. This filter is much wider than any configuration tested in our work, either in simulation or in experiment, but as the simulations demonstrate, for pulses longer than 400 ps FWHM (300 ps FWHM at 1 cm SDS, supplement Figure S2), the CNR for the cerebral signal is reduced. Further the constraint of using a time-of-flight filter with the shape of a Bessel function, a consequence of the modulation, introduces additional limitations due to the side lobes of the coherence gate, which may allow for short pathlength interference. While careful tuning of the width and position of the time-of-flight filter was performed to reduce the influence of short pathlength light, the linking of gate center time and gate width is undesirable. This link is not required in our approach that uses a nearly transform-limited, pulsed laser, which enables both a gating-function duration within the optimal range for human cerebral blood flow measurements as well as independent selection for the gate center time. Methods utilizing much shorter coherence lengths have also been demonstrated by Safi et al. [38] and Zhang et al. [39]. In these works, the coherence gate width FWHM was ∼1.9 ps and ∼39 ps, respectively. Extrapolating the results of this work, shortening the pulse duration beyond the range that was tested would also lead to a decrease in the CNR for the cerebral blood flow signal. The reported width of the coherence gate, source-detector separation used, minimum autocorrelation temporal lag achievable, number of independent speckle measurements, BF_i_ sampling rate, and the centroid of the filtered time of flight distribution relative to the peak of the TPSF for each technique is summarized in Table 2. The use of an optimal coherence gate width allows for the selection of a gate center that both allows for sufficient SNR of the measurement and a sufficiently late distribution of time-of-flights. While methods based on coherence gating have been predominately discussed and compared to PaLS-iDCS, other methods have been developed to measure pathlength-specific light intensity and are based on low coherence interferometry [54,55], non-linear wavelength up-conversion gating [56–58], and higher order, multi-frequency correlation functions [59]. Although valuable, these are not well suited for the specific requirements of cerebral blood flow monitoring, namely the ability to measure μs-scale fluctuations in light intensity at late times of flight (>500ps after the peak of the TPSF) with sufficient SNR to enable temporal sampling of BF_i_ at a rate where the features of the cardiac pulsation can be resolved.

**Table 2.** Comparison of interferometric, pathlength-selective, diffuse optical methods.

| | Coherence gate width (FWHM, ps) | Source-detector separation (cm) | Minimum autocorrelation lag (μs) | Number of independent speckle measurements | Sampling rate (Hz) | Approximate centroid of the latest arriving filtered time of flight distribution relative to the peak of the TPSF (ps) |
| --- | --- | --- | --- | --- | --- | --- |
| Safi et al. [38] | 1.9 | 2.5 | 1 | 1 | 20 | 460 |
| Zhang et al. [39] | 39 | Transmission | 294 | 1 | 1.7 | 127 |
| This work | 360 | 2 | 1 | 1 | 20 | 730 |
| Zhao et al. [37] | 1600 | 1 | 3 | $\sim 200$ | 100 | 450 |

The implementation of pathlength selection demonstrated here has the potential to provide optimal cerebral blood flow measurements, and, as is the case for CW-iDCS systems, is additionally able to scale to multichannel systems [30,31]. The ability to convert a reference pathlength sweep into an optical property measurement allows for quantification of optical properties without needing to acquire the TPSF using TCSPC hardware, enabling more accurate estimation of the BF_i_ when the autocorrelation function is fit with the proper optical properties. This functionality was shown to be consistent across liquid phantom experiments across pulse durations and in healthy subject experiments, showing its robustness in the face of changing experimental conditions used to optimize the CNR of the cerebral blood flow measurement. As a limitation of the current instrumentation, the pathlength sweep did take a relatively long time to complete (∼30 minutes), and variability of the absorption and scattering was observed in the simultaneously collected TD-DCS measurements (shown in supplemental section S7). The sweep time could be improved, for example, by using a fiber optic switch with preset delays, enabling a faster sweep through a sequence of pre-selected reference arm lengths and allowing for a more consistent optical property measurement. Though single photon detectors and TCSPC electronics were used in this work to thoroughly characterize the technique, this is not required for PaLS-iDCS. Non-photon counting detectors can sustain much higher sample arm signal levels, allowing the distance between source and detector to be reduced, and also side-stepping the hold-off time and pile-up effects that limit the overall count rate of single photon measurements. While helped by the 2 cm source-detector separation, photon count rate limitations didn’t allow us to experimentally overwhelm the entire sample arm with the reference arm and we used TCSPC for proper gate selection to select photons in a ToF gate where the reference arm was much stronger than the gated sample arm. This limitation can be removed by using a non-photon counting detector, enabling both the ability to measure at shorter source-detector separation, and increasing the absolute number of deep photons (supplemental section S4) as compared to longer separations. Further, the use of camera-type sensors would give us the ability to employ highly parallelized detection, increasing averaging and SNR, allowing for greatly simplified instrumentation [30] – to be explored in future work. To conclude, PaLS-iDCS has the potential to advance both functional imaging as well as clinical cerebral blood flow monitoring by providing signals that are both high SNR and have high sensitivity to brain physiology.

## Data Availability

The data that support the findings of this study are available from the corresponding author on reasonable request.

## Supplementary content

See supplement 1 for additional supporting information.

