## Supplemental material for "Pathlength-selective, interferometric diffuse correlation spectroscopy"

### S1. Derivation of the coherence length based on the interference between delayed, transform limited, gaussian pulses

We define two gaussian pulses that are split from the same laser, which is assumed to have phase stability throughout the duration of the pulse, i.e. the electric field of the source at a time, t, is given by an oscillating term multiplied by an envelope $\left(E(t) = \exp(j\omega t) * \exp\left(-\frac{t^2}{4\sigma^2}\right)\right)$. The two pulses propagate along different paths and are combined at a detector. We express the electric field at the detector as,

$$E(t) = E_1(t, x_1) + E_2(t, x_2), \tag{S.1}$$

where $E_1(t, x_1)$ and $E_2(t, x_2)$ are the representations for the divided pulse as a function of time, t, and pathlength distance, $x_i$. The individual electric field of the pulse is given as,

$$E_i(t, x_i) = \exp(j\omega t)\exp(-jkx_i)\exp\left(-\frac{\left(t - \frac{x_i}{\nu}\right)^2}{4\sigma^2}\right), \tag{S.2}$$

where $\omega$ is the frequency of the light $\left(\omega = \frac{c_0}{\lambda_0}\right)$, k is the wavenumber of the light in the medium $\left(k = \frac{2\pi n}{\lambda_0}\right)$, $\nu$ is the speed of light in the medium $\left(\nu = \frac{c_0}{n}\right)$, and $\sigma$ is the temporal standard deviation of the intensity pulse where the full width at half maximum (FWHM) of the intensity pulse is given by $FWHM = 2\sqrt{2\log(2)}\,\sigma$. The intensity measured at the detector is given as,

$$I(t) = \exp\left(-\frac{\left(t - \frac{x_1}{\nu}\right)^2}{2\sigma^2}\right) + \exp\left(-\frac{\left(t - \frac{x_2}{\nu}\right)^2}{2\sigma^2}\right) + \exp\left(-\frac{\left(t - \frac{x_1}{\nu}\right)^2 + \left(t - \frac{x_2}{\nu}\right)^2}{4\sigma^2}\right)\left(2\cos\left(k(x_1 - x_2)\right)\right). \tag{S.3}$$

We can define an average pathlength, $\bar{x} = \frac{x_1 + x_2}{2}$, and the pathlength difference, $\Delta x = x_1 - x_2$, which allows for the simplification of the interference term. Substituting the mean pathlength and pathlength difference expressions into Equation S.3 gives,

$$I(t) = \exp\left(-\frac{\left(t-\frac{x_1}{v}\right)^2}{2\sigma^2}\right) + \exp\left(-\frac{\left(t-\frac{x_2}{v}\right)^2}{2\sigma^2}\right) + \exp\left(-\frac{\left(t-\frac{\bar{x}}{v}\right)^2}{2\sigma^2}\right)\exp\left(-\frac{\Delta x^2}{8v^2\sigma^2}\right)(2\cos(k\Delta x)). \quad (S.4)$$

For a slow detector that cannot resolve the temporal shape of the pulse, the intensity detected will be proportional to the integral of the intensity with respect to time, given as

$$I(\Delta x) = 2\sigma\sqrt{2\pi} + 2\sigma\sqrt{2\pi}\exp\left(-\frac{\Delta x^2}{8v^2\sigma^2}\right)\cos(k\Delta x). \quad (S.5)$$

The coherence length, $l_c$, is defined as the pathlength difference, Δx, at which the amplitude of the interference term normalized by the value of the intensity at a large pathlength difference ($\Delta x \to \infty$) falls to 1/e (~37%)[1]. With this definition, we can define the coherence length in terms of the pulse duration, given in Equation S.6,

$$l_c = \frac{FWHM}{\sqrt{\log(2)}}v. \quad (S.6)$$

This derived relationship connecting the pulse duration to the coherence length is used for the simulations.

### S2. Derivation of the form of the PaLS-iDCS autocorrelation function

The measured autocorrelation functions from the PaLS-iDCS instrument can be described similarly to the form of autocorrelation functions collected by a time-domain DCS instrument[2] with the addition of the terms describing the reference arm. The electric field impinging on the detector is described in Equation S.7 as a sum of the sample electric field and the reference electric field,

$$E(t_M, s) = E_S(t_M, s) + E_R(t_M, s), \quad (S.7)$$

where $E_S(t,s)$ and $E_R(t,s)$ are the electric field terms for the sample and reference, respectively, at a macroscale experimental time, $t_M$, and electric field trajectory pathlength, *s*. We describe the form of the sample arm electric field term in Equation S.8 as a sum of *N* independent trajectories at each trajectory pathlength, *s*,

$$E_S(t_M, s) = \sum_{i=0}^{N} E_{S,i}(t_M, s) = \sqrt{\frac{\langle I_S(s)\rangle}{N}}\exp\left(j\left(\frac{2\pi v}{\lambda}t_M - \frac{2\pi}{\lambda}s\right)\right)\sum_{i=1}^{N}\exp\left(-j\Delta\phi_{s,i}(t_M)\right), (S.8)$$

where $\langle I_S(s)\rangle$ is the average detected intensity for a trajectory of length *s*, *v* is the speed of light, $\lambda$ is the wavelength of light, and $\Delta\phi_{s,i}$ represents the difference in phase for the individual emitter, *i*, arising from dynamic scattering events along the trajectories. The total detected intensity from the sample arm is described as the integral of the average intensity across all trajectory pathlengths, i.e. $\langle I_S\rangle = \int_0^\infty \langle I_S(s)\rangle ds$, where the pathlength distribution of the light intensity, *P(s)*, is given by the ratio of the average intensity at a given pathlength to the total average intensity, i.e. $\langle I_S(s)\rangle/\langle I_S\rangle$. The collected pathlength distribution reflects the tissue optical absorption and scattering properties as governed by light propagation in the diffuse regime. For a measurement of a semi-infinite media in the reflectance geometry, by applying the method of images to satisfy the boundary condition [3], the remitted pathlength distribution can be described as Equation S.9,

$$P(s) = \left(\sqrt{\frac{4\pi}{3\mu_s' r_1^2}}\exp\left(-\sqrt{3\mu_a\mu_s' r_1^2}\right) - \sqrt{\frac{4\pi}{3\mu_s' r_2^2}}\exp\left(-\sqrt{3\mu_a\mu_s' r_2^2}\right)\right)^{-1} \cdot s^{-\frac{3}{2}}\left(\exp\left(-\sqrt{\frac{3\mu_s' r_1^2}{4s}}\right) - \exp\left(-\sqrt{\frac{3\mu_s' r_2^2}{4s}}\right)\right)\exp(-\mu_a s), \quad (S.9)$$

where $\mu_s'$ is the reduced scattering coefficient; $\mu_a$ is the optical absorption coefficient; $r_1$ and $r_2$ are the distances from the sources in the method of images to the detector and are described as $r_1 = \sqrt{\rho^2 + \frac{1}{\mu_s'^2}}$ and $r_2 = \sqrt{\rho^2 + \left(\frac{1}{\mu_s'} + 2z_b\right)^2}$ ; ρ is the source-detector separation; $z_b$ is the extrapolation distance to the imaging boundary and is computed as $z_b = \frac{2}{3\mu_s'}\left(\frac{1+R_{eff}(n)}{1-R_{eff}(n)}\right)$; and $R_{eff}$ accounts for the mismatch in index of refraction between the tissue and the air and is estimated as a function of the index of refraction, given as $R_{eff}(n) = -1.44n^{-2} + 0.71n^{-1} + 0.668 + 0.0636n$. The reference arm electric field is modeled as a single trajectory with a pathlength $l_R$, given as $E_R(t_M, s) = \sqrt{\langle I_R\rangle}\exp\left(j\left(\frac{2\pi v}{\lambda}t_M - \frac{2\pi}{\lambda}s\right)\right)\delta(s - l_R)$. The measured light intensity is described in Equation S.10,

$$I(t_M, s) = E_S(t_M, s)E_S^*(t_M, s) + E_R(t_M, s)E_R^*(t_M, s) + E_S(t_M, s)E_R^*(t_M, s) + E_S^*(t_M, s)E_R(t_M, s), \quad (S.10)$$

which is a sum of the intensity of the sample arm, $E_S(t_M, s)E_S^*(t_M, s)$, at a given macroscale time, $t_M$, and pathlength, $s$, the intensity of the reference arm, $E_R(t_M, s)E_R^*(t_M, s)$, and the conjugate pair of interference terms between the sample and reference electric fields. With simplification for terms where the expected value is equal to zero or where there is a product of temporally uncorrelated non-zero terms, the expanded form of the unnormalized intensity autocorrelation function as a function of trajectory pathlength, $G_2(s, \tau)$, is given in Equation S.11,

$$\begin{aligned} G_2(s,\tau) = &\langle I_S(t_M, s)I_S(t_M + \tau, s)\rangle + 2\langle I_S(s)\rangle\langle I_R(s)\rangle + \langle I_R(t_M, s)I_R(t_M + \tau, s)\rangle \\ &+ \langle E_S(t_M, s)E_R^*(t_M, s)E_S^*(t_M + \tau, s)E_R(t_M + \tau, s)\rangle \\ &+ \langle E_S^*(t_M, s)E_R(t_M, s)E_S(t_M + \tau, s)E_R^*(t_M + \tau, s)\rangle. \end{aligned} \quad (S.11)$$

This expression for the unnormalized intensity autocorrelation function represents an ideal case, where instrumentation effects like the instrument response function (IRF), the mutual coherence of trajectories with different pathlengths ($\Gamma(\Delta s)$), and the number of detected modes are not included. Following the derivation provided in Cheng et al [2], we express the full, unnormalized, PaLS-iDCS intensity autocorrelation function as a function of experimental gate selection time, $t_g$, in Equation S.12,

$$\begin{aligned} G_2(t_g, \tau) = &\iint I_S(s)IRF\left(t_g - \frac{s}{v}\right)I_S(s')IRF\left(t_g - \frac{s'}{v}\right)ds\,ds' \\ &+ 2\iint I_S(s)IRF\left(t_g - \frac{s}{v}\right)I_R(s')IRF\left(t_g - \frac{s'}{v}\right)ds\,ds' \\ &+ \iint I_R(s)IRF\left(t_g - \frac{s}{v}\right)I_R(s')IRF\left(t_g - \frac{s'}{v}\right)ds\,ds' \\ &+ \frac{\beta_p}{M}\iint I_S(s)IRF\left(t_g - \frac{s}{v}\right)g_{1,S}(\tau, s)I_S(s')IRF\left(t_g - \frac{s'}{v}\right)g_{1,S}(\tau, s')\Gamma(s - s')ds\,ds' \\ &+ \frac{2\beta_p}{M}\iint I_S(s)IRF\left(t_g - \frac{s}{v}\right)g_{1,S}(\tau, s)I_R(s')IRF\left(t_g - \frac{s'}{v}\right)g_{1,R}(\tau, s')\Gamma(s - s')ds\,ds', \end{aligned} \quad (S.12)$$

where M is the number of detected spatial modes, $\beta_p$ is the polarization-dependent coherence factor, computed as $\beta_p = \frac{I_\parallel^2 + I_\perp^2}{(I_\parallel + I_\perp)^2}$, where $I_\parallel$ and $I_\perp$ represent the intensity carried by two orthogonal polarization components, and $g_{1,S}(\tau, s)$ and $g_{1,R}(\tau, s)$ are the pathlength specific, normalized electric field temporal autocorrelation functions for the sample arm and reference arm, respectively. The first three terms of Equation S.12 represent the baseline value of the autocorrelation (i.e. as $\tau \to \infty$), the fourth term is the standard TD-DCS autocorrelation that was previously derived, and the fifth term is autocorrelation function arising from the interference between sample and reference arms. The mutual coherence function for a source with a gaussian spectral profile is defined as $\Gamma(\Delta s) = \exp\left(-\frac{2\Delta s^2}{l_c^2}\right)$, where $l_c$ is the coherence length of the source, where the definition given in Section S.1 is used. To simplify the expression in Equation S.12, we will define several terms to condense the expression. An effective pathlength distribution describing the blurring caused by the IRF is given as $P_X'(s, t_g) = P_X(s) IRF\left(t_g - \frac{s}{v}\right)$. We define the fraction of the collected intensity, $f_X(t_g)$, in either the sample or reference arm at a particular gate selection time, as the integral of the effective pathlength distribution, i.e. $f_X(t_g) = \int_0^\infty P_X'(s, t_g) ds$. We can also see that the form of the reference arm terms can be utilized to simplify the last term of Equation S.12. The normalized electric field autocorrelation of the reference arm, $g_{1,R}(\tau, s)$, can be computed from the assumed form of the reference arm electric field and is found to be equal to 1 for all time lags, τ. Further, due to the pathlength distribution of the reference arm being a delta function, the integration in the last term can be greatly simplified. To normalize the form of the correlation function, we compute the square of the mean total intensity collected in the IRF gate, given in Equation S.13,

$$\begin{aligned} \langle I_T(t_g)\rangle^2 &= \left\langle \int I_S(t_M, s) IRF\left(t_g - \frac{s}{v}\right) ds + \int I_R(t_M, s) IRF\left(t_g - \frac{s}{v}\right) ds \right\rangle^2 \\ &= \langle I_S\rangle^2 f_S(t_s)^2 + 2\langle I_S\rangle f_S(t_s)\langle I_R\rangle f_S(t_R) + \langle I_R\rangle^2 f_R(t_s)^2. \end{aligned} \qquad (S.13)$$

Applying each of these simplifications, we can compute the normalized intensity autocorrelation function, $g_2(t_g, \tau)$, given in Equation S.14,

$$\begin{aligned} g_2(t_g, \tau) = 1 + \frac{\beta_p}{M\langle I_T(t_g)\rangle^2} \Bigg( &\langle I_S\rangle^2 \iint P_S'(s, t_g) g_{1,S}(\tau, s) P_S'(s', t_g) g_{1,S}(\tau, s') \exp\left(-\frac{2(s - s')^2}{l_c^2}\right) ds ds' \\ &+ 2\langle I_S\rangle\langle I_R\rangle \int P_S'(s, t_g) g_{1,S}(\tau, s) IRF\left(t_g - \frac{l_R}{v}\right) \exp\left(-\frac{2(s - l_R)^2}{l_c^2}\right) ds \Bigg). \end{aligned} \qquad (S.14)$$

Based on Equation S.14, given we measure the reference intensity, we can also estimate the sample intensity at a given gate selection time by evaluating the form of the overall coherence factor, $\beta(t_g)$, which is defined as the value of the normalized autocorrelation function at zero lag ($\tau = 0$) minus 1. Given in Equation S.15, the form of the overall coherence parameter is,

$$\begin{aligned} \beta(t_g) = \frac{\beta_p}{M\langle I_T(t_g)\rangle^2} \Bigg( &\langle I_S\rangle^2 \iint P_S'(s, t_g) P_S'(s', t_g) \exp\left(-\frac{2(s - s')^2}{l_c^2}\right) ds ds' \\ &+ 2\langle I_S\rangle\langle I_R\rangle \int P_S'(s, t_g) IRF\left(t_g - \frac{l_R}{v}\right) \exp\left(-\frac{2(s - l_R)^2}{l_c^2}\right) ds \Bigg). \end{aligned} \qquad (S.15)$$

While the term proportional to $\langle I_S\rangle^2$ in the parentheses in Equations S.14 and S.15 is included for completeness, in the typical conditions where $\langle I_R\rangle \gg \langle I_S\rangle$, it will be negligible. Equations S.14 and S.15 are restated in the main text as Equations 5 and 6 with reference to the derivation provided here.

### S3. Measurement of the coherence properties of the pulsed laser source

While for the simulations we can reasonably assume a gaussian profile for wavelength spectrum of the laser source, for our pulsed laser, while the shaping done allows for the creation of a pulse shape closer to that of a transform-limited pulse, the coherence properties are not perfectly described by the simplified gaussian model. To estimate the coherence properties of the laser source at the different pulse durations, we constructed a Mach-Zehnder interferometer to measure the fringe contrast at different pathlength differences. To accomplish this, the 99% output of the 99%/1% fused fiber coupler connected to the output of the optical amplifier (Fig 1) was coupled into a 50%/50% fused fiber coupler (TN1064R5A2A, Thorlabs) and split into two arms. One arm was coupled directly to a collimating lens (F220APC-1064, Thorlabs), while the other was sent through the variable length delay line used in the full system (Fig 6). The delayed arm was coupled into a collimating lens (F220APC-1064, Thorlabs), and the two collimated beams were superimposed at a small angle on a CMOS camera (acA1300-200um, Basler) and the fringe contrast was estimated at different path offsets in steps of 1 cm. Example fringe images are seen in Fig S1.a, and the fringe amplitude across the central ROI of the images are modeled using the expression given in Equation S.16.

$$I(x,y) = I_1(x,y) + I_2(x,y) + 2\sqrt{I_1(x,y)I_2(x,y)}\cos\big(\phi_1(x,y) - \phi_2(x,y)\big)\,\gamma^{(1)}(\Delta l) \quad (S.16)$$

where $I_1$ and $I_2$ are the spatial distributions on the axes of the camera (x,y) of the intensity from the two beams of the interferometer, $\phi_1$ and $\phi_2$ are the spatial distributions of the phase of the two beams, and $\gamma^{(1)}$ is the degree of coherence between the beams at a pathlength difference $\Delta l$. To estimate the degree of coherence between the two beams at a particular pathlength difference, four images were taken: (1) dark, (2) $I_1$, (3) $I_2$, and (4) $I_1+I_2$. From these four images, we computed the magnitude of the interference term by subtracting the dark image from images 2, 3 and 4, then subtracting images 2 and 3 from image 4. The final scaling was to divide image 4 by 2 times the square root of the product of images 2 and 3. This leaves the sinusoidal interference fringes scaled by the degree of coherence. By estimating the normalized amplitude of the fringes, the degree of coherence was estimated for different pathlength differences, $\Delta l$. To estimate any coherence loss due to the pulsed seed laser, we seeded the same laser shaping optical circuit with a long-coherence CW laser (RFLM-1064nm, NP Photonics) and measured the coherence length of the resulting output. Finally, we estimate the coherence length of the laser as though the pulsed amplitude were an envelope for a perfect sinusoidal signal with no amplitude-phase coupling introduced by the shaping of the pulse by the EOM. We find that for the pulse shape generated by our pulser board, the coherence length relative to a gaussian pulse ($l_c$ = 10.8 cm) with the same FWHM duration is 2.7 cm shorter for the optimal, chirpless case ($l_c$ = 8.1 cm), 3.8 cm shorter for the shaping of the long-coherence source ($l_c$ = 7.0 cm), and 4.5 cm shorter for the shaping of the pulsed laser ($l_c$ = 6.3 cm) (Fig S1.c). In Figure S1.b, the normalized fringe contrast for three example pathlength offsets are shown for the shaped, pulsed seed laser.

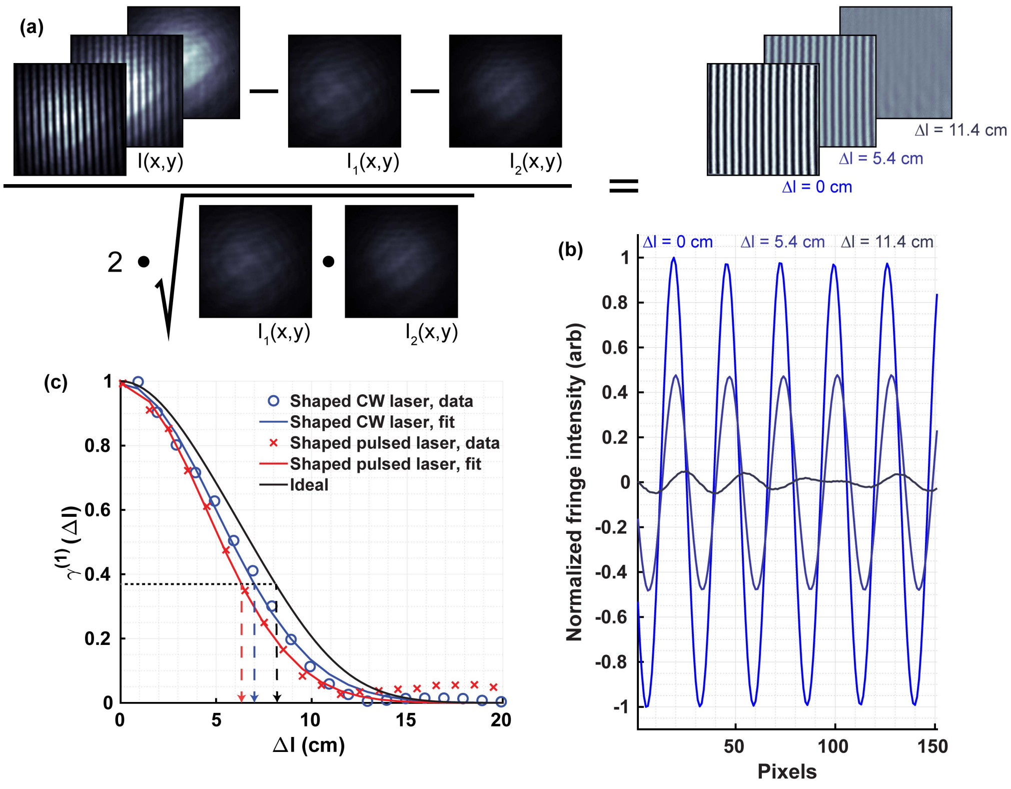


**Figure S1:** (a) The image processing performed to estimate the mutual coherence function for the shaped laser pulse is shown for three example offsets of 0 cm, 5.4 cm, and 11.4 cm. (b) The normalized fringe contrast is averaged across the camera and plotted for the three example pathlength offsets. (c) Comparison of the value of the mutual coherence function for the shaped, pulsed laser ($l_c$ = 6.3 cm), the shaped, CW laser ($l_c$ = 7.0 cm), and the ideal envelope case ($l_c$ = 8.1 cm) for the 300 ps FWHM pulse shape. A gaussian pulse with the same FWHM has a coherence length of 10.8 cm.

### S4. Comparison of simulated measurements made at shorter source-detector separations

In the main text, the simulations presented maintain the same source-detector separation used in both the phantom and in-vivo experiments. The use of a 2 cm source-detector separation was selected to keep the photon detection rate in check. This is needed to minimize the effects of non-linearity introduced by the single photon detector hold-off time, which limits the maximal overall count rate of the detector, and the pile-up effect, which reduces the probability of detecting photons after the peak of the TPSF. For detection systems that do not require TCSPC acquisition and single photon detectors, so long as the detector is not saturated and the light from the reference arm greatly exceeds the return from the sample, measurements can be made at shorter source-detector separations, which will increase the overall number of photons that have traveled to the brain, allowing for further enhanced measurements[4]. To characterize this improvement, simulations were performed for a measurement taken at a 1 cm source-detector separation, and the results are quantified in the same manner as was done for the results presented in Figure 1 in the main text and shown in Figure S2 below. Improvement in the maximal achievable CNR is seen for both TD-DCS and PaLS-iDCS (Fig S2.b, S2.e, and S2.h), showing further improvement over the performance of the CW-iDCS technique at the range of simulated source-detector separations between 5 mm and 40 mm. While the TD-DCS performance is much improved relative to the 2 cm measurement as well as in comparison to the PaLS-iDCS measurement at 1 cm, due to the higher expected count rate at the 1 cm source-detector separation (~10x that at 2 cm), with current single photon counting detector technology suited for TD-DCS, significant distortion of the shape of the TPSF and autocorrelation functions would be expected, leading to inaccurate measurements of tissue optical properties and cerebral blood flow. Because PaLS-iDCS does not require single photon counting detectors, these limitations are removed, and even higher SNR can be achieved from the interferometric technique.

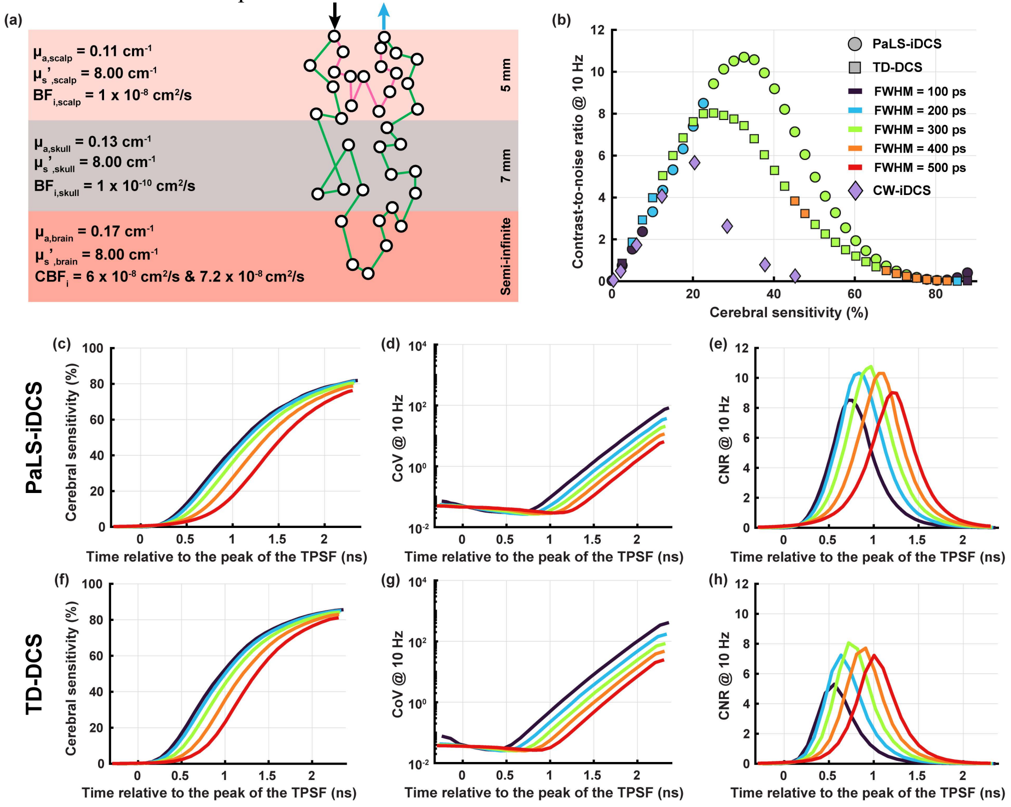


**Figure S2:** (a) Depiction of the simulation geometry with tissue layer optical properties. The photon trajectories demonstrate the shortened source-detector separation, and the relative shape of the sensitivity profile of the measurement. In (b) the CNR of the simulated measurements are compared as a function of the sensitivity of the measurement to the cerebral layer. For both TD-DCS and PaLS-iDCS, the maximal CNR and the sensitivity at the maximal CNR are both increased relative to the values reached for the 2 cm measurement. The pulse duration providing the highest CNR at each sensitivity level is also shifted toward shorter pulse durations, likely due to the interaction between the narrower pathlength distribution that is observed at shorter source-detector separations and the blurring of blood flow information that comes with longer pulse durations. The CW-iDCS results are presented again as a reference for comparison. For the shorter source-detector separation measurement, for the same time gate after the peak of the TPSF, the sensitivity (c and f) is reduced relative to the 2 cm measurement, though the same maximal sensitivity is reached by going to longer times of flight. The shorter source-detector separation greatly increases the number of photons collected at later time gates, and this is reflected in (d and g), where the CoV of the measurement remains much lower for time gates much later in the time-of-flight distribution. Combining the sensitivity and COV gives the figures shown in (e and h), which demonstrate both improved CNR relative to the longer source-detector separation measurement and the shift of the optimal pulse duration to a shorter FWHM.

### S5. Effect of gate duration on the simulated contrast-to-noise ratio of measured $BF_i$ of TD-DCS

In the main text, for both simulated and measured data, the presented results were analyzed using a gate duration of 5/3 the FWHM of the pulse, as previous work has demonstrated improvements to measurement SNR were seen to plateau at this gate duration[5]. To expand on the analysis presented in the main text and compare the effect gate duration has on the expected contrast-to-noise ratio for the cerebral signal, for the TD-DCS simulation data, we compute the three gate durations specified in Table 1 for each pulse duration and plot the estimated CNR vs. the cerebral sensitivity of the measurement in Figure S3. For all pulse durations, the selection of the intermediate pulse duration (5/3 FWHM) provides the highest achievable CNR as well as higher CNR at higher cerebral sensitivities as compared to either the longer or shorter gate duration. The use of too short of a photon selection gate, while providing a more pathlength-specific autocorrelation function, results in a reduced number of photon arrivals being used to compute the autocorrelation function which directly decreases the SNR of the measurement. For too long a photon selection gate, a greater number of photon detections happening near the peak of the TPSF are typically included in the gate, except for gates with exceedingly late gate center times, which increases the influence of the scalp hemodynamics on the measured signal. This reduces the sensitivity of the measurement to cerebral hemodynamics, though due to the form of the noise of the autocorrelation function, the additional photons collected do not appreciably decrease the coefficient of variation of the fit $BF_i$, and the result is an overall reduction in measurement CNR. By selecting the intermediate gate, a balance is struck between the inclusion of sufficient photon number to maintain SNR while also maintaining pathlength specificity.

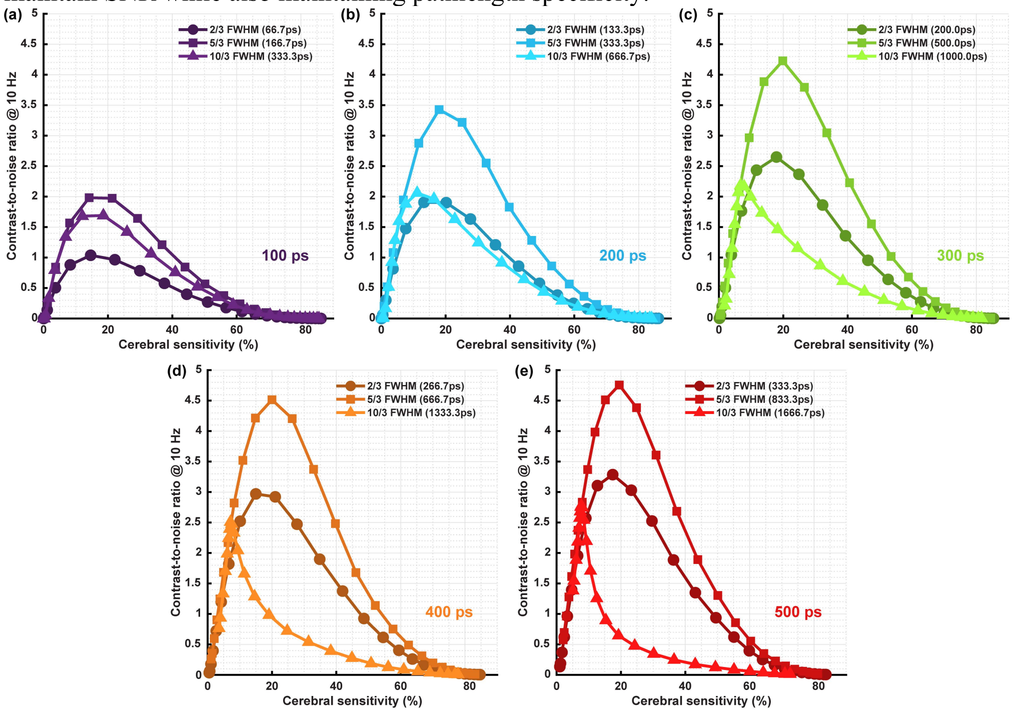


**Figure S3.** Comparison of the TD-DCS contrast-to-noise ratio as a function of cerebral sensitivity for pulse durations of (a) 100 ps FWHM, (b) 200 ps FWHM, (c) 300 ps FWHM, (d) 400 ps FWHM, and (e) 500 ps FWHM. For all pulse durations, the intermediate gate duration provided the highest achieved CNR as well as increased CNR at higher values of cerebral sensitivity.

### S6. Comparison of unnormalized reduction in relative $BF_i$ and relative $pBF_i$ during the pressure modulation task

In the main text, for the comparison of both steady state and pulsatile $BF_i$ metrics during the pressure modulation maneuver (Figure 5), the results were presented normalized by the drop in the short separation $BF_i$ signals to address the possible inconsistencies in the applied pressure and its effect on the blood flow. In Figure S4.a and S4.b, we plot the distribution of relative changes for each individual channel (i.e. 5 mm, early gate, and late gate) without normalization by the relative change in the short separation signal. For both steady state and pulsatile metrics, as in the main text, the reduction in $BF_i$ is less at the later gate, indicating a reduced sensitivity to the extracerebral hemodynamics, though the distinction between responses is reduced due to the experimental variability.

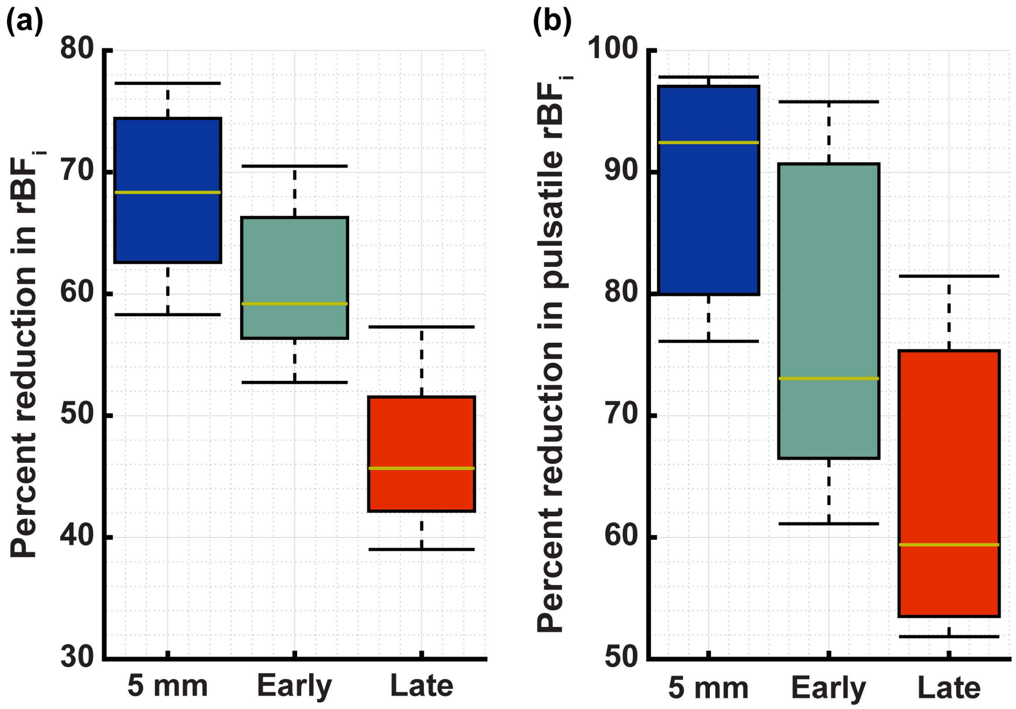


**Figure S4.** Comparison of the reductions in steady state (a.) and pulsatile (b.) relative $BF_i$ during the pressure modulation task for the 5 mm short separation, the early gate, and the late gate. The increase in the spread of the distributions as compared to the results presented in Figure 5 in the main text demonstrates the effect of the experimental variability (i.e. different pressure modulation pressure, different extracerebral thicknesses between subjects, etc.). The unnormalized changes still allow for the distinction of the three different signals, indicating a robust increase in cerebral sensitivity afforded by the late gate.

### S7. Optical property variability during the reference arm pathlength sweep

Due to the relatively long duration of the reference arm pathlength sweep used to assess the optical absorption and scattering with PaLS-iDCS in the human subject measurements, changes in the optical properties during the measurement interval may disrupt the accurate recovery of the optical properties. To assess variability over the measurement interval, for each subject, optical properties are estimated at each reference arm position from the simultaneously measured TD-DCS channel. In Figure S.5, the changes relative to the mean absorption (S.5a) and mean reduced scattering (S.5b) coefficients are shown as a function of the reference arm position for the PaLS-iDCS measurement. A linear model is fit for all subject data for both optical properties and the fit line as well as the 99% confidence interval is overlaid over the individual data points. An overall increase in both absorption and scattering is observed from the aggregated subject data, though this change is relatively small in magnitude (±0.0025 $cm^{-1}$ in $\mu_a$, ±0.225 $cm^{-1}$ in $\mu_s$'), representing less than a 7% relative change in both properties relative to the mean value presented in Figure 4.d. While a small change, faster pathlength sweeps would further reduce the range of optical properties sampled and improve the accuracy of the measurement.

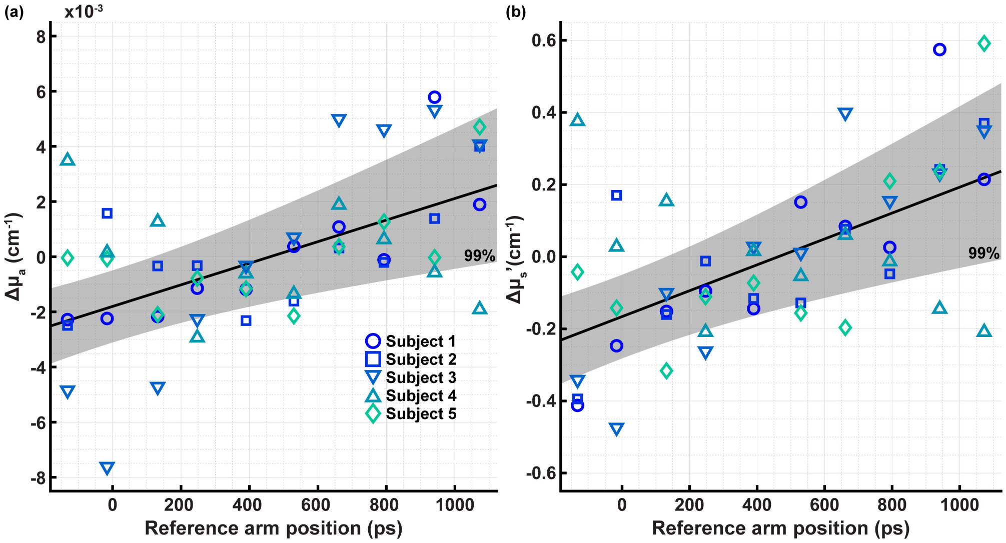


**Figure S5.** Comparison of the changes in absorption (a.) and reduced scattering (b.) measured with TD-DCS at each timepoint corresponding to an individual reference arm position. Variability in the optical properties as a function of reference arm position were observed and could be contributing factors to the observed inaccuracies in the estimated optical properties seen in the main text. Increasing the speed of the reference arm sweep would allow for a more accurate assessment of the tissue optical properties by sampling the properties in a period of time where they remain more stable.
